# A Promoter Competition Hub Orchestrates *Runx1* Alternative Promoter Usage during Skeletal Muscle Stem Cell Activation

**DOI:** 10.64898/2026.07.13.738242

**Authors:** Liangqiang He, Qiang Sun, Yulong Qiao, Ziliu Wang, Qin Zhou, Lifang Han, Zhenguo Wu, Hao Sun, Huating Wang

## Abstract

Alternative promoter (AP) usage profoundly expands transcriptomic and proteomic diversity, yet the regulatory principles governing promoter choice remain poorly understood. Here, using the dual-promoter *Runx1* locus as a paradigm in skeletal muscle stem cells (MuSCs), we uncover a promoter competition hub that orchestrates AP selection during MuSC fate transition from activation to proliferation. We find antagonistic expression dynamics between the two *Runx1* promoters, with the primary promoter (PP) active in the early activating stage and the secondary promoter (SP) induced later in proliferation. Combinatorial genetic perturbations *in vitro* and *in vivo* establish the non-redundant roles of PP- and SP-derived isoforms in MuSC lineage progression. Mechanistically, we find that PP and SP engage in reciprocal competition in a multi-connected enhancer-promoter (E-P) loop hub. Further dissection identifies a cohort of key enhancers that dynamically interact with PP/SP to orchestrate the competition. Moreover, we establish the transcription factor USF1 as a key factor driving the dynamic E-P interactions and PP/SP competition. Beyond *Runx1*, high-resolution Micro-C reveals promoter competition as a general mechanism regulating a subset of AP choices in MuSC activation/proliferation. Collectively, our study uncovers a dynamic promoter competition hub that governs AP selection in MuSC lineage progression, offering fundamental insights into how 3D genome architecture coordinates precise, stage-specific gene expression programs during cell fate transitions.

## Introduction

Promoters are central determinants of transcriptional output. Accumulated studies have shown that most human and mouse protein-coding genes are controlled by multiple promoters^1–3^. The use of different promoters, known as alternative promoters (APs), generates mRNA isoforms with varying 5′-untranslated region (UTR) lengths, substantially increasing transcript diversity and also influencing post-transcriptional processes^1,3^. In some cases, AP-derived isoforms can encode functionally distinct protein products, further expanding proteomic complexity and functional repertoire^1,4^. Utilization of APs is widespread across diverse species, and their dysregulation has been increasingly linked to human diseases. For example, integrative analyses of cell-fate transitions in yeast have uncovered a sophisticated regulatory network governing AP usage, which is critical for the dynamic fine-tuning of gene expression^4^. Pan-cancer transcriptomic profiling also demonstrates pervasive promoter dysregulation across tissues, cancer subtypes, and individual patients, with promoter activity emerging as a more robust prognostic marker for patient survival than overall gene expression^3^. Moreover, recent studies have revealed coordinated crosstalk between transcription start and end sites, demonstrating that the choice of 5′ end AP directly influences 3′ end selection^5,6^. Several mechanisms have been implicated in directing AP usage. It has been shown that DNA methylation, histone modifications, and chromatin remodeling at promoter regions are key determinants of AP selection^4,7–9^. Notably, aberrant DNA methylation can drive the AP usage with a significant negative correlation observed between methylation levels and promoter activity^9^. Additionally, the binding of cell-type specific transcription factors (TFs) can also play a role in the process of AP choice^10,11^. Nevertheless, the in-depth mechanistic dissection of the AP usage remains incomplete. We believe elucidating such mechanisms will shed novel insights into broader principles governing promoter choice.

Gene expression is precisely governed by the intricate communication between promoters and their cognate enhancers, with dysregulation of these interactions leading to aberrant transcriptional profiles^12–15^. A central unresolved question in gene regulation is how distal enhancers selectively transmit regulatory signals to their target promoters. 3D genome organization has emerged as an important mechanism of enhancer-promoter (E-P) interactions. Topologically associating domains (TADs) can mediate enhancer selectivity by promoting E-P interactions within the same TAD and by preventing E-P interactions between different TADs^12,13,15^. The architectural protein CTCF is thought to play an important role by functioning as an E-P insulator and/or E-P facilitator, other factors including cohesin, Mediator, even transcriptional machinery and Pol II can also facilitate E-P interactions^12,13,15^. In addition, some TFs are actively involved in mediating E-P communication through different mechanisms including direct protein dimerization/oligomerization, interactions with architectural factors, or biomolecular condensation and phase separation^13^. Moreover, emerging evidence suggests that local promoter architecture, including promoter-specific motifs, GC content, and CpG islands, also dictates the compatibility, stability, and efficiency of E-P interactions^15,16^. The complication has been escalated by recent studies revealing the formation of E-P hubs within which either one promoter interacts with multiple different enhancers (promoter hubs) or one enhancer connects to multiple promoters (enhancer hubs)^13,17^; multi-connected E-P hubs, on the other hand, refer to hubs with multiple promoters interacting with multiple enhancers. The multi-connected E-P hubs are thought to play an important role in allowing regulatory information to be transmitted to multiple genes to allow coordinated changes in gene expression during cell-state transitions^13,17,18^. Nevertheless, their organizational principles and mechanisms of assembly remain poorly understood. We reason that AP usage could represent a unique scenario for dissecting E-P interaction and multi-connected E-P hub organization.

In this study, we elucidated the mechanisms governing AP usage and promoter choice on the paradigm of the dual-promoter *Runx1* locus in the setting of skeletal muscle stem cell (MuSC) fate transition during acute injury-induced muscle regeneration. MuSCs are indispensable for skeletal muscle regeneration and homeostasis^19^. In the quiescent state, MuSCs are characterized by high expression of the TF *Pax7*. Upon injury, quiescent MuSCs (QSCs) rapidly activate to form myoblasts (activated MuSCs, ASCs), marked by the induction of basic helix-loop-helix (bHLH) TFs *Myf5* and *MyoD*. Most ASCs downregulate *Pax7* expression and undergo myogenic differentiation to form new muscle fibers, while a subpopulation of ASCs with high *Pax7* expression returns to quiescence to replenish the MuSC pool. TF-driven intrinsic gene regulatory network governing the MuSC fate transitions, especially in the phase of activation, is a focused area of research^20–22^. RUNX1 (Runt-related transcription factor 1) is upregulated following injury and its deletion in MuSCs impairs proliferation and muscle repair^23,24^. RUNX1 is well studied for its essential function in hematopoiesis^25^ and known to possess two well-defined promoters driving its transcription: the proximal P2 and the distal P1^26^. In addition, transcripts from these two promoters undergo alternative splicing to generate multiple distinct protein isoforms that significantly expand the complexity of the *Runx1* repertoire^27,28^. The precise spatiotemporal control of P1/P2 promoter usage and the resulting balance of RUNX1 isoforms are critical for fine-tuning hematopoietic stem cell emergence, progenitor expansion, and lineage differentiation^27–29^. Genetic ablation of P1 impairs the initial emergence of hematopoietic progenitors and disrupts lineage fidelity in adult hematopoiesis, whereas attenuated P2 function causes severe hematopoietic failure and midgestational embryonic lethality^30^. However, the underlying mechanism of P1 vs. P2 choice remains a puzzle, representing a prime paradigm for dissecting AP usage and promoter choice.

Here in this study, we found that AP usage of *Runx1* locus is also present in MuSC fate transitions during muscle regeneration with primary promoter (PP, known as P2)- and secondary promoter (SP, known as P1)-derived *Runx1* transcripts exhibiting opposing expression dynamics. Moreover, the derived protein isoforms exert distinct roles during MuSC activation process. Further investigation integrating multi-omics analyses and combinatorial genetic perturbation assays demonstrates an antagonism of the two promoters within a multiway E-P hub. Moreover, we identified a cohort of key enhancers involved in the E-P rewiring and PP/SP competition during MuSC activation. Interestingly, we found that CTCF is dispensable for *Runx1* promoter competition. USF1 on the other hand binds to the SP, modulates E-P connectivity within the hub and drives isoform-specific transcription, thereby representing a key TF orchestrating MuSC fate transition and muscle regeneration. Lastly, extending beyond the *Runx1* paradigm, we found AP usage is a general phenomenon occurring at many loci during MuSC activation and promoter competition represents a general mechanism regulating a subset of AP usage.

## Results

### Temporal dynamics of *Runx1* alternative promoter usage during MuSC fate transitions in myogenic lineage progression

To elucidate the *Runx1* AP usage during MuSC fate transitions, we leveraged our previously published^31^ and in-house transcriptomic profiles of MuSCs at distinct lineage stages (Fig. 1a, Extended Data Fig. 1 and Supplementary Table 1). Briefly, freshly isolated MuSCs (FISCs) (considered as early activating MuSCs^32,33^) were collected from *Pax7-nGFP* mice via fluorescence-activated cell sorting (FACS) and cultured *in vitro* for 1, 2, or 3 days to generate activated MuSCs (ASC-D1), proliferating myoblasts (ASC-D2), or early differentiating myocytes (DSC-D3). Quiescent MuSCs (QSCs) were acquired through *in situ* paraformaldehyde (PFA) fixation prior to FACS isolation, a method known to preserve MuSC quiescence^32,33^. We examined the expression dynamics of *Runx1* mRNA isoforms derived from P1 and P2, respectively (Fig. 1b) and found that the two promoters exhibited distinct temporal expression patterns during the lineage progression (Fig. 1b-1c). While both transcripts were barely detectable in QSCs (FPKM: 3.86 for P1 and 5.66 for P2), P2-derived isoforms were dramatically induced in FISCs (FPKM: 112.63) followed by progressive decline in ASC-D1 (FPKM: 50.35), ASC-D2 (FPKM: 19.71) and DSC-D3 (FPKM:17.61). Conversely, P1 was not induced until D1 (FPKM: 1.06 in FISCs and 2.42 in D1) and continued to elevate its expression (FPKM: 19.34 and 34.45 in D2 and D3), revealing a clear shift in promoter usage from P2 to P1 when MuSCs progressed along the myogenic lineage. We therefore designated P2 as the primary promoter (PP) and P1 as the secondary promoter (SP). Consistent with the above transcriptomic profiles, ChIP-seq for H3K4me3, a marker of active transcription, detected peaks exclusively at the PP region in FISC, whereas both SP and PP regions showed prominent peaks in ASC-D2 (Fig. 1b and Supplementary Table 2). Similarly, ATAC-seq and H3K27ac signals were detected on PP in both FISC and ASC-D2, whereas SP exhibited similar chromatin features exclusively in ASC-D2. To further validate the above discovered *Runx1* AP usage dynamics, we designed SP- and PP-specific primers for qRT-PCR quantification of the corresponding *Runx1* isoforms (Fig. 1d). As expected, the expression of SP-derived isoforms (*SP-Runx1*) increased along with *MyoD* expression, whereas *PP-Runx1* decreased along with the *Pax7* expression (Fig. 1e). Accordingly, the total RUNX1 protein level gradually escalated during MuSC activation, with multiple bands of varying sizes suggesting the possible presence of distinct protein isoforms derived from the two promoters (Fig. 1f).

**Figure 1.**
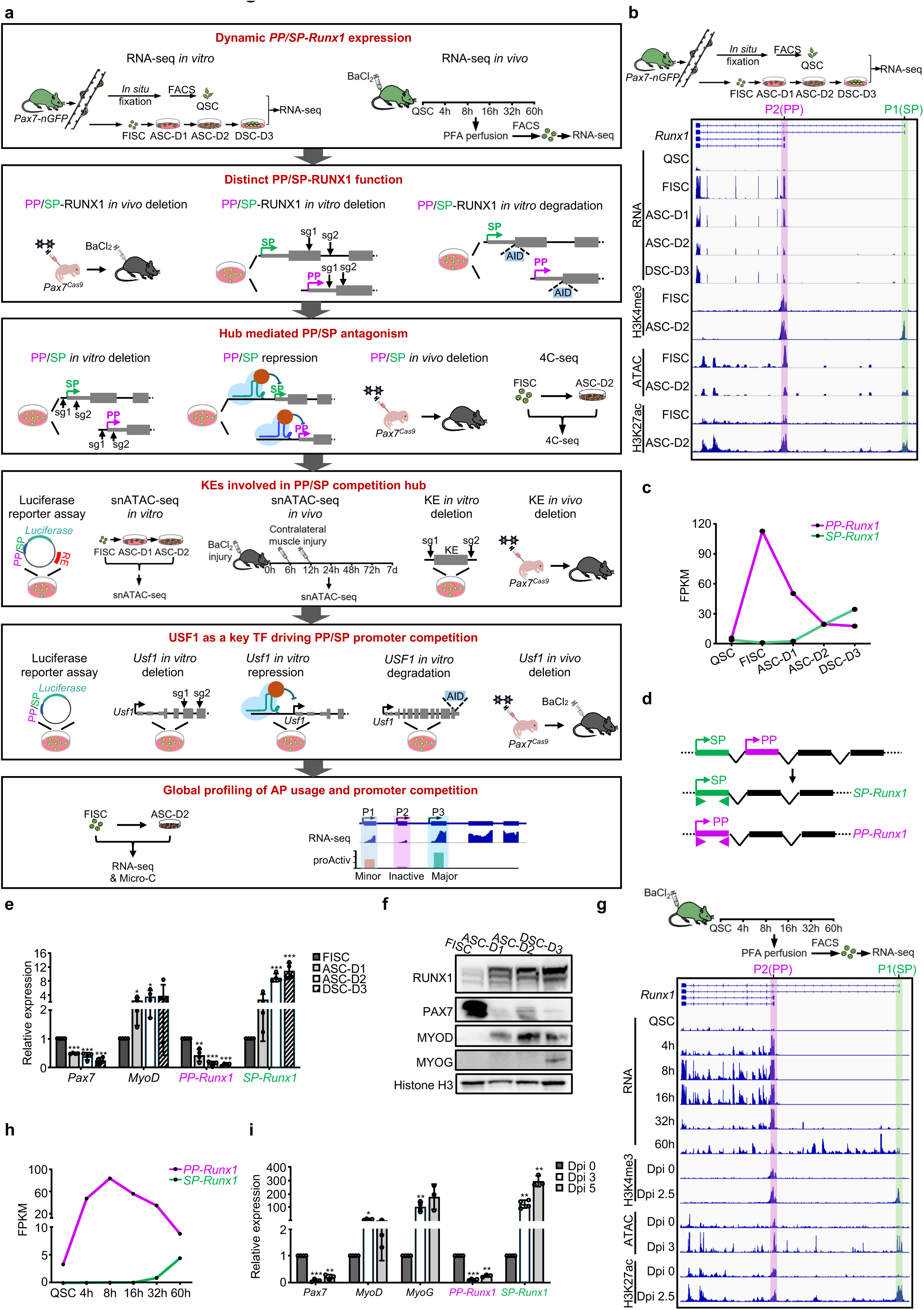
Temporal dynamics of *Runx1* alternative promoter usage during MuSC fate transitions in myogenic lineage progression. (**a**) Schematic of the overall design of the study. **(b)** RNA-seq tracks showing the expression dynamics of *PP-* and *SP-Runx1* isoforms during MuSC *in vitro* lineage progression. ChIP-seq profiling of H3K4me3 and H3K27ac along with ATAC-seq signals at the *Runx1* locus in FISCs and ASC-D2 is also presented. Schematic of MuSC collection from *Pax7-nGFP* mice and the *in vitro* lineage progression is shown above. **(c)** Th expression FPKM values of *PP*- and *SP-Runx1* transcripts from the above RNA-seq data is shown. FPKM: fragments per kilobase per million mapped reads. (**d**) Schematic illustration of the qRT-PCR primers designed for specific detection of *PP-* and *SP-Runx1* isoforms. (**e**) qRT-PCR analysis showing the expression dynamics of *PP-* and *SP-Runx1*. (**f**) Western blot of RUNX1, PAX7, MYOD, MYOG protein levels in the above MuSCs. Histone H3 was used as a loading control. (**g**) RNA-seq tracks depicting the expression dynamics of *PP-* and *SP-Runx1* isoforms in MuSCs collected from the *in vivo* lineage progression. ChIP-seq for H3K4me3 and H3K27ac as well as ATAC-seq signals at the *Runx1* locus at Dpi 0 and 2.5-3 are also shown. Schematic of MuSC collection from the BaCl_2_ injury-induced regenerating muscle at the indicated time points is shown above. (**h**) The expression FPKM values of *PP*- and *SP-Runx1* transcripts from the above RNA-seq data is shown. (**i**) qRT-PCR analysis of *PP-* and *SP-Runx1* expression in MuSCs isolated at Dpi 0, 3, and 5. Statistical significance was calculated using Student’s t-test for (**e**) and (**i**). *p < 0.05, **p < 0.01, ***p < 0.001.

To substantiate the above observed *Runx1* promoter switch, we examined the expression of PP/SP-derived *Runx1* isoforms in MuSCs *in vivo* in acute injury induced regenerating muscle. BaCl_2_ was injected into the tibialis anterior (TA) muscle to induce acute injury, which resulted in extensive myofiber necrosis, followed by sequential phases of immune cell infiltration and MuSC activation within hours post-injury^34^. Activated MuSCs subsequently proliferated and differentiated into new myofibers leading to muscle repair within three to four weeks post-injury^34^. Accordingly, we analyzed publicly available RNA-seq data from *in situ* fixed MuSCs isolated at various time points after BaCl_2_-induced muscle injury^35^ (Fig. 1g-1h). Mirroring the *in vitro* findings, *PP-Runx1* expression was drastically induced 4-8 hours post-injury when MuSCs began to activate, and subsequently declined progressively to 60 hours as MuSCs underwent proliferation and differentiation. *SP-Runx1* expression was not detected until 32 hours post-injury and continued to increase at 60 hours. Correspondingly, the PP locus exhibited prominent H3K4me3, H3K27ac ChIP-seq, and ATAC-seq signals in MuSCs isolated from both 0 (corresponding to FISCs) and 2.5-3 days post injury (Dpi) (corresponding to ASCs). In contrast, the SP locus displayed such peaks only in MuSCs at Dpi 2.5-3 (Fig. 1g), recapitulating the *in vitro* dynamics (Fig. 1b). The above results were confirmed by qRT-PCR analysis of MuSCs collected from Dpi 0, 3 and 5, which revealed a progressive increase in *SP-Runx1* expression. In contrast, the level of *PP-Runx1* was markedly downregulated from Dpi 0 to 3, with a slight recovery observed by Dpi 5 (Fig. 1i). Collectively, the above results demonstrate the temporal dynamics of *Runx1* AP usage during MuSC fate transitions in myogenic lineage progression and a clear switch from PP to SP during cell activation.

### Distinct roles of PP- and SP-derived isoforms in MuSC activation/proliferation

Given the distinct expression dynamics of PP- and SP-derived mRNA isoforms during MuSC activation process (Fig. 1c and 1h), we speculated that the derived proteins may play different roles with PP-RUNX1 promoting MuSC activation while SP-RUNX1 arising later to facilitate myoblast proliferation. Notably, since alternative splicing also occurs among the *Runx1* isoforms generated from both PP and SP^27,28^, PP-RUNX1 and SP-RUNX1 collectively refer to all protein isoforms originating from each promoter, respectively. To test our speculation, we perturbed their expression in QSCs using previously established muscle-specific *in vivo* CRISPR/Cas9/AAV9-sgRNA editing system^36,37^(Fig. 2a). Briefly, AAV9-sgRNAs targeting *PP*- or *SP-Runx1* specific coding regions (4 × 10^11^ viral genomes (vg) per mouse) were intramuscularly (IM) injected into *Pax7^Cas^*^9^ mice at postnatal day 5 (P5), followed by a second injection (8 × 10^11^ vg per mouse) at P10 (Fig. 2a). Control mice received AAV9-sgRNA vector backbone without any sgRNA insertion. Targeted deep sequencing revealed efficientj editing of the PP and SP coding regions in MuSCs from the treated mice (71.4% and 64.7% editing efficiency respectively) (Fig. 2b) accompanied by substantial reduction of total RUNX1 protein levels (Fig. 2c). MuSCs from the mice were isolated and analyzed for their activation/proliferation. As expected, perturbation of PP-RUNX1 impaired MuSC activation (Fig. 2d); retarded proliferation was also observed probably as a consequence of impaired activation (Fig. 2e); SP-RUNX1 deficiency however had no impact on MuSC activation (Fig. 2f) but significantly retarded their proliferative capacity (Fig. 2g).

**Figure 2.**
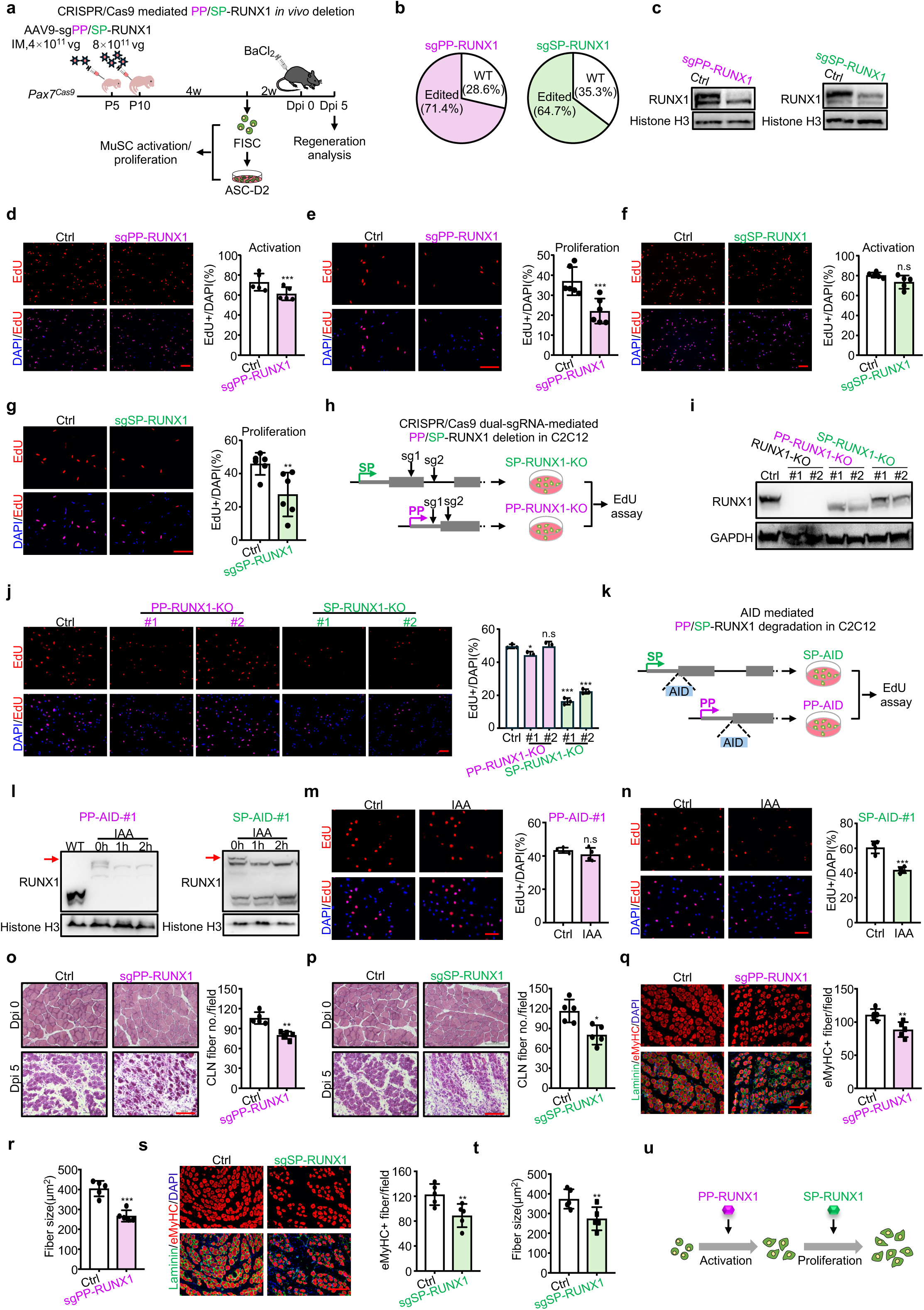
Distinct roles of PP- and SP-derived isoforms in MuSC activation/proliferation. (**a**) Schematic of *in vivo Runx1* isoform-specific knockout in MuSCs. AAV9 vectors encoding sgRNAs targeting PP- or SP-specific coding regions (4 × 10¹¹ vg/mouse at P5, followed by 8 × 10¹¹ vg/mouse at P10) (sgPP or sgSP) were injected intramuscularly into *Pax7^Cas9^* mice. Control mice received the same dose of AAV9 virus containing the pAAV9-sgRNA backbone without sgRNA insertion. FISCs were harvested and cultured to ASC-D2 cells for analysis. For muscle regeneration assays, BaCl₂ was injected into the TA muscle at least six weeks post-AAV injection (Dpi 0), and samples were collected five days later (Dpi 5) for analysis. (**b**) The above MuSCs were isolated and subjected to targeted deep sequencing to assess editing efficiency at sgRNA targeted regions. (**c**) Western blot showing RUNX1 protein expression in MuSCs isolated from the above PP-RUNX1 (Left) or SP-RUNX1 (Right) knockout mice. Histone H3 served as a loading control. (**d**) Left: FISCs from the Ctrl or sgPP injected mice were isolated and labeled with EdU for 24 hours to assess MuSC activation. Right: Quantification of the percentage of EdU+ cells. Scale bar = 100 μm. (**e**) Left: ASCs from the above mice were labeled with EdU to evaluate MuSC proliferation. Right: Quantification of the percentage of EdU+ cells. Scale bar = 100 μm. (**f**) Left: FISCs from the with Ctrl or sgSP injected mice were isolated and labeled with EdU for 24 hours to assess MuSC activation. Right: Quantification of the percentage of EdU+ cells. Scale bar = 100 μm. (**g**) Left: ASCs from the above mice were labeled with EdU to evaluate MuSC proliferation. Right: Quantification of the percentage of EdU+ cells. Scale bar = 100 μm. (**h**) Schematic illustration for generating C2C12 cells with PP-RUNX1 or SP-RUNX1 knockout using dual sgRNA mediated CRISPR/Cas9 deletion. (**i**) Western blot showing RUNX1 protein expression in the above generated SP-RUNX1-KO, PP-RUNX1-KO, and total RUNX1-KO C2C12 cells. GAPDH served as loading control. (**j**) Left: EdU staining of Ctrl, PP-RUNX1-KO, and SP-RUNX1-KO C2C12 cells. Right: Quantification of the percentage of EdU+ cells. Scale bar = 100 μm. (**k**) Schematic for generating C2C12 cell lines with inducible PP-RUNX1 or SP-RUNX1 degradation using the AID2 system. The mAID tag was inserted at N-terminus of PP-RUNX1 or SP-RUNX1 protein. (**l**) Western blot showing degradation of PP-RUNX1 and SP-RUNX1 following treatment of PP-AID-#1 and SP-AID-#1 clones with 5-Ph-IAA (IAA). The degraded protein bands are indicated with red arrows. Histone H3 was used as a loading control. (**m**) Left: EdU staining of the above IAA treated clone PP-AID-#1cells. Right: Quantification of EdU+ cells. Scale bar: 100 μm. (**n**) Left: EdU staining of the above IAA treated clone SP-AID-#1 cells. Right: Quantification of EdU+ cells. Scale bar: 100 μm. (**o**-**p**) Left: H&E staining of TA muscles at Dpi 5 from the sgPP-RUNX1 (**o**) or sgSP-RUNX1(**p**) mice generated in (**a**). Right: Quantification of regenerating myofibers with CLN per field. Scale bar: 100 μm. (**q**) Left: Immunostaining of eMyHC (Red) and Laminin (Green) on the above TA muscles from sgPP-RUNX1 mice. Right: Quantification of eMyHC+ myofibers. Scale bar: 100 μm. **(r)** Quantification of myofiber size. (**s**) Left: Immunostaining of eMyHC (Red) and Laminin (Green) on the TA sections from sgSP-RUNX1 mice. Right: Quantification of eMyHC+ myofibers. Scale bar: 100 μm. (**t**) Quantification of myofiber size. (**u**) Schematic illustration of differential roles of PP- or SP-RUNX1 proteins in MuSCs. Statistical significance was calculated using Student’s t-test. *p < 0.05, **p < 0.01, ***p < 0.001 and ns, no significance.

To substantiate the finding, we also generated C2C12 myoblast cell lines with specifical knockout of PP- or SP-RUNX1 (coding regions were edited) by CRISPR/Cas9-mediated genome editing (Fig. 2h and Extended Data Fig. 2a-2b) to assess the impact on myoblast proliferation. Total RUNX1 knockout C2C12 cells were also generated alongside, in which no detectable RUNX1 protein was observed (Fig. 2i). Deletion of PP-RUNX1 primarily abolished the upper RUNX1 protein isoform, while loss of SP-RUNX1 led to a global reduction in total RUNX1 protein levels (Fig. 2i), comparable to results observed *in vivo* (Fig. 2c). Functional assessment by EdU incorporation assays revealed that PP-RUNX1-KO had no obvious effect, whereas SP-RUNX1-KO significantly impaired C2C12 proliferation (Fig. 2j). Furthermore, we generated C2C12 myoblast cell line with inducible degradation of PP- or SP-RUNX1 using the auxin-inducible degron (AID2) technology^38,39^. A mAID tag was inserted at the N-terminus of either PP- or SP-RUNX1 in C2C12 cells stably expressing the engineered auxin receptor OsTIR1 (F74G) (Fig. 2k and Extended Data Fig. 2c). The addition of 5-Ph-IAA (auxin, IAA) induced rapid degradation of a portion of the total RUNX1 protein in both clones (Fig. 2l and Extended Data Fig. 2d). Consistently, PP-RUNX1 deletion had no obvious impact (Fig. 2m and Extended Data Fig. 2e), whereas loss of SP-RUNX1 (Fig. 2n and Extended Data Fig. 2f) profoundly impaired myoblast proliferation.

To further dissect the functional importance of PP/SP-RUNX1, we examined their effects on muscle regeneration (Fig. 2a). Six weeks after AAV-sgRNA virus injection, BaCl₂-induced muscle injury was performed, and muscle regeneration was assessed at Dpi 5. PP- and SP-RUNX1 deficiency both resulted in a reduced number of newly regenerating myofibers characterized by centrally localized nuclei (CLN) (Fig. 2o-2p). Immunofluorescence (IF) staining for embryonic myosin heavy chain (eMyHC) also confirmed a marked decrease in the number and size of the newly formed myofibers (Fig. 2q-2t), underscoring the essential roles of both isoforms in acute injury-induced muscle regeneration. Collectively, our results reveal distinct and stage-specific roles for PP- and SP-derived isoforms: *PP-Runx1* is induced early and promotes MuSC activation, while *SP-Runx1* is turned on later to facilitate myoblast proliferation (Fig. 2u).

### *Runx1* PP/SP promoter competition orchestrated by E-P interaction rewiring in a multi-connected E-P hub

Next, we aimed to explore the mechanism underlying *Runx1* PP/SP switching during MuSC activation. The mutually exclusive usage dynamics suggested a likely promoter competition model. To test the notion, we specifically deleted PP or SP (promoters only, excluding the gene body sequences) in C2C12 cells (Fig. 3a and Extended Data Fig. 3a). As expected, knockout of PP (PP-KO) significantly attenuated *PP-Runx1* expression accompanied by elevated *SP-Runx1* expression (Fig. 3b, left); a reciprocal pattern was observed upon SP deletion (SP-KO) (Fig. 3b, right), supporting the idea of promoter competition. To further validate the notion, we employed CRISPR/dCas9-KRAB-mediated transcriptional repression (CRISPRi) to generate PP- or SP-Krab C2C12 cells in which PP or SP was silenced respectively (Fig. 3c). As expected, reduced *PP-Runx1* expression was observed in PP-Krab cells accompanied by increased *SP-Runx1* expression (Fig. 3d, left); analogous results were detected in SP-Krab cells (Fig. 3d, right). Expectedly, PP repression also reduced Pol II binding, H3K4me3 and H3K27ac modifications at PP but correspondingly elevated levels were observed at SP (Fig. 3e, top and Extended Data Fig. 3b); similar reciprocal changes were observed upon SP repression (Fig. 3e, bottom). Finally, we tested whether PP/SP competition exists during MuSC activation *in vivo*. To this end, PP or SP was deleted *in vivo* by the CRISPR/Cas9/AAV9-sgRNA system (Fig. 3f and Extended Data Fig. 3c). PP deletion in sgPP mice resulted in a robust and sustained decrease of *PP-Runx1* across FISC, D1, and D2 stages, with a corresponding increase of *SP-Runx1* levels at all time points (Fig. 3g and Extended Data Fig. 3d). In contrast, SP deletion in sgSP mice significantly diminished *SP-Runx1* level only at D2 (Fig. 3h, right and Extended Data Fig. 3e, right), which corresponded to elevated *PP-Runx1* level (Fig. 3h, left and Extended Data Fig. 3e, left). Altogether the above genetic evidence supports a promoter competition model that drives the mutually exclusive expression dynamics of PP and SP in MuSCs.

**Figure 3.**
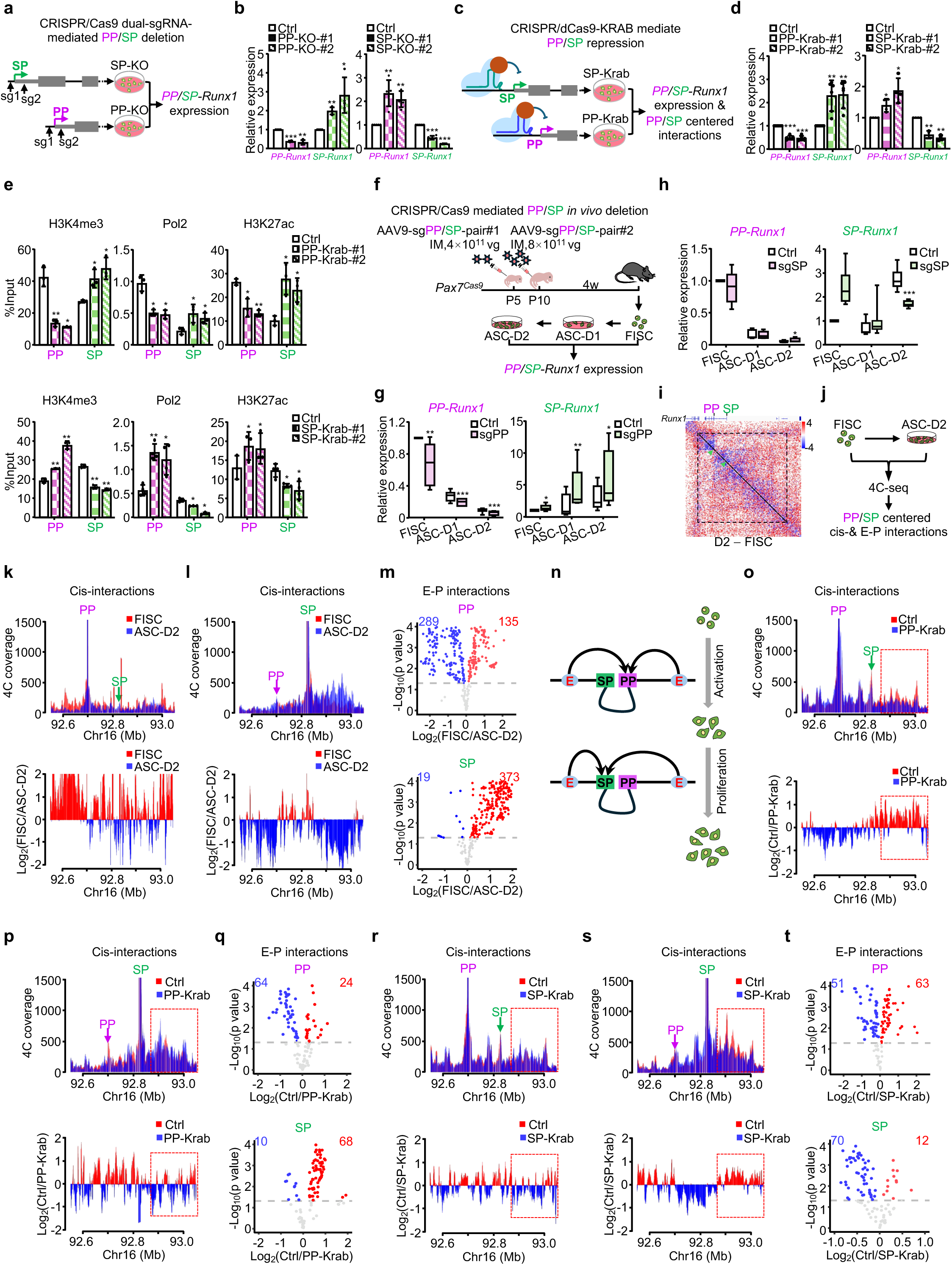
*Runx1* PP/SP promoter competition orchestrated by E-P interaction rewiring in a multi-connected E-P hub. (**a**) Schematic illustration showing the design of generating PP-or SP knockout C2C12 myoblast cells by dual sgRNAs mediated targeted deletion. (**b**) qRT-PCR detection of *PP-* and *SP-Runx1* expression in the above cells. (**c**) Schematic illustration of dCas9-KRAB-mediated repression of PP or SP. (**d**) qRT-PCR detection of *PP-* and *SP-Runx1* expression in the above cells. (**e**) ChIP-qPCR analysis of H3K4me3 (Left), RNA Pol II (Middle), and H3K27ac (Right) occupancy at the PP and SP in the above PP (Top)- or SP (Bottom)-KRAB cells. (**f**) Schematic of *in vivo* knockout of PP or SP in MuSCs. *Pax7^Cas9^*mice were intramuscularly injected with 4 × 10¹¹ vg/mouse of AAV9-sgRNA pair #1 at P5 and followed by 8 × 10¹¹ vg/mouse of AAV9-sgRNA pair #2 at P10 targeting PP or SP. Control mice received the same dose of AAV9 virus containing the pAAV9-sgRNA backbone without sgRNA insertion. FISCs were isolated four weeks later and cultured to ASC-D1 and ASC-D2 to detect *PP-* and *SP-Runx1* expression by qRT-PCR. (**g**-**h**) qRT-PCR detection of *PP-* (Left) and *SP-Runx1* (Right) expressions in the above sgPP(**g**) or sgSP (**h**) cells. (**j**) Heatmap of existing Hi-C data showing differential contact map at genomic regions encompassing the *Runx1* locus in FISCs and ASC-D2. The TAD harboring *Runx1* is indicated by a dashed triangular. The locations of PP and SP in the heatmap are marked with green arrows. (**k**) Schematic illustrating the experimental workflow of the 4C-seq to detect PP/SP centered cis- and E-P interactions in FISC and ASC-D2. (**k**-**l**) 4C-seq analysis showing the change of PP/SP centered cis-interactions in FISC and ASC-D2. Top: Normalized 4C-seq coverage profiles showing overlaid contact signals at PP (**k**) or SP (**l**) locus in FISC (Red) vs. ASC-D2 (Blue). The reads are normalized to 1M total reads. Bottom: Differential cis-interactions centered on PP (**k**)- or SP (**l**) between FISC and ASC-D2. (**m**) The number of increased (Red) or decreased (Blue) E-P interactions at PP (Top) and SP (Bottom) in FISC vs. ASC-D2. (**n**) Schematic illustration of PP and SP competition during MuSC activation and proliferation. (**o**-**p**) 4C-seq analysis showing the change of PP/SP centered cis-interactions in Ctrl and PP-Krab cells. Top: Normalized 4C-seq coverage profiles showing overlaid contact signals at PP (**o**) or SP (**p**) in Ctrl (Red) vs. PP-Krab (Blue) cells. The reads are normalized to 1M total reads. Bottom: Differential cis-interactions centered on PP (**o**)- or SP (**p**) between Ctrl and PP-Krab cells. The region upstream of the SP is highlighted with a red rectangle. (**q**) The number of increased (Red) or decreased (Blue) E-P interactions at PP (Top) and SP (Bottom) in Ctrl vs. PP-Krab. (**r**-**s**) 4C-seq analysis showing the change of PP/SP centered cis-interactions in Ctrl and SP-Krab cells. Top: Normalized 4C-seq coverage profiles showing overlaid contact signals at PP (**r**) or SP (**s**) in Ctrl (Red) vs. SP-Krab (Blue) cells. The reads are normalized to 1M total reads. Bottom: Differential cis-interactions centered on PP (**r**)- or SP (**s**) between Ctrl and SP-Krab cells. The region upstream of the SP is highlighted with a red rectangle. (**t**) The number of increased (Red) or decreased (Blue) E-P interactions at PP (Top) and SP (Bottom) in Ctrl vs. SP-Krab. Statistical significance was calculated using Student’s t-test. *p < 0.05, **p < 0.01, ***p < 0.001 and ns, no significance.

Given the well-established role of chromatin interaction in promoter regulation, we hypothesized that the PP/SP competition might be driven by rewiring of chromatin interactions. Comparison of our existing Hi-C data^33^ in FISC and ASC-D2 revealed that the *Runx1* locus resided within a distinct TAD (Fig. 3i and Extended Data Fig. 3f). Expectedly, PP-centered chromatin interactions were markedly attenuated during MuSC activation (Fig. 3i); in contrast, changes in SP-centered interactions were not obvious (Fig. 3i). Further examination by 4C-seq (Fig. 3j) confirmed that as cells progressed from FISC to ASC-D2, PP-centered total cis-interaction number diminished (Fig. 3k and Extended Data Fig. 3g, top), while SP-centered cis-interactions increased (Fig. 3l and Extended Data Fig. 3g, bottom). Specifically, on PP, 135 E-P interactions were gained or strengthened, while 289 were lost or diminished (Fig. 3m, top). On SP, 373 E-P interactions were gained or enhanced and only 19 lost or weakened (Fig. 3m, bottom). Interestingly, direct interactions between PP and SP were also detected (Fig. 3k-3l). Altogether the above observations suggest that PP and SP may collectively form a 3D interaction hub, within which PP and SP compete for enhancers leading to mutual repression (Fig. 3n). Supporting our speculation, we found that silencing PP in the PP-Krab cells significantly reduced PP-centered total cis-interactions (Fig. 3o and Extended Data Fig. 3h, top), particularly within the intergenic regions upstream of SP, while SP-centered interactions were concomitantly enhanced (Fig. 3p and Extended Data Fig. 3h, bottom). Consistently, 24 E-P interactions were gained or strengthened on PP, while 64 were lost or diminished (Fig. 3q, top). Conversely, SP displayed 68 gained or enhanced E-P interactions, with only 10 being lost or weakened (3q, bottom). Repression of SP in the SP-Krab cells resulted in similar but reciprocal changes in PP- and SP-centered total cis- (Fig. 3r-3s and Extended Data Fig. 3i) and E-P (Fig. 3t) interactions. Again, interactions between PP and SP were detected (Fig. 3o-3p and 3r-3s), suggesting a persistent physical association despite functional antagonism. Collectively, results from the above integrated analyses support that SP and PP engage in reciprocal competition possibly in a multi-connected E-P loop hub to achieve their distinct temporal expression dynamics during MuSC activation.

### Identification of regulatory hub elements orchestrating *Runx1* PP/SP competition

To further dissect the hub architecture governing *Runx1* promoter choice, we sought to identify key enhancers that form the E-P hub and mediate the PP/SP competition. By integrating H3K27ac ChIP-seq, ATAC-seq, and publicly available key TF (PAX7, MYOD, MYOG, c-JUN) binding profiles from FISCs and ASCs, we defined a total of 24 candidate regulatory elements (RE1-RE24) within the *Runx1* locus (Fig. 4a). Among them, 14 (RE1-RE14) resided in the intergenic regions upstream of SP and exhibited prominent H3K27ac signals especially at the ASC stage, whereas the remaining 10 (RE15-RE24) were located within *Runx1* introns, with a subset (RE21-RE24) showing obvious H3K27ac enrichment in both FISCs and ASCs. These REs were also enriched for key TF binding. Specifically, PAX7 was observed on RE1, RE6, RE22 and RE23, while c-JUN, MYOD and MYOG binding was present at most of the REs. Luciferase assays using reporters in which each RE was fused to PP or SP revealed that most of the REs displayed varying degrees of enhancer activity toward PP (Fig. 4b, left). In contrast, only a subset exhibited significant enhancer activity for SP (Fig. 4b, right). Notably, seven REs, including RE1, RE6, RE11, RE12, RE13, RE14, and RE22, demonstrated robust enhancer activity on both promoters thus defined as key enhancers (KEs). Further analysis of the 4C-seq data revealed that interactions between all these KEs and SP increased during MuSC activation (Fig. 4c, bottom), consistent with the observed upregulation of SP-centered chromatin contacts (Fig. 3l and 3m). The interactions with PP were however not all decreased: while KE1, KE11, KE12, KE22 showed diminished interactions with PP, KE6 and KE14 exhibited increased interactions and KE13 remained unchanged (Fig. 4c, top); this is consistent with the markedly diminished but sustained PP expression at D2 (Fig. 1c).

**Figure 4.**
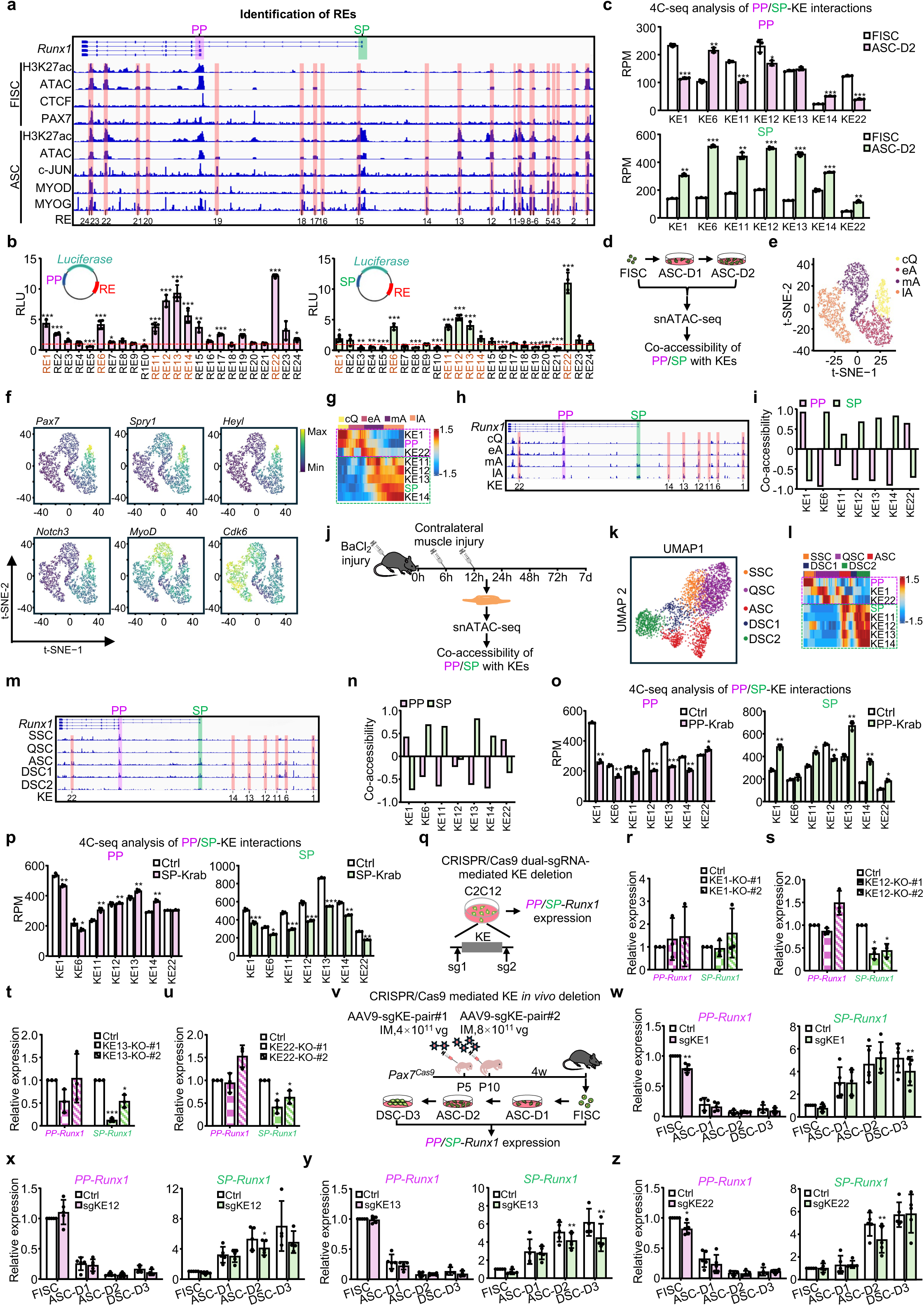
Identification of regulatory hub elements orchestrating *Runx1* PP/SP competition. (**a**) Identification of 24 REs (RE1 to RE24) in the genomic region surrounding the *Runx1* locus. (**b**) Luciferase reporter assay to assess the enhancer activity of the identified REs in ASC-D2 using either PP (Left) or SP (Right) as the minimal promoter. The constructed luciferase reporter plasmids are shown. RLU: relative luciferase unit. The red dashed line indicates an RLU of 1. (**c**) 4C-seq analysis showing interactions between the indicated KEs and PP (top) or SP (bottom) in FISC and ASC-D2. RPM: reads per million total reads. (**d**) Schematic illustrating the experimental workflow of the snATAC-seq assay performed in FISC, ASC-D1, and ASC-D2. (**e**) t-SNE clustering showing four sub-clusters of MuSCs. (**f**) t-SNE plots displaying the gene scores of the indicated genes across the four sub-clusters. (**g**) Heatmap showing the chromatin accessibility dynamics of the PP/SP and the KEs along the pseudotime trajectory inferred from the snATAC-seq data. Green and purple rectangles highlight KEs with accessibility patterns co-varying with PP and SP, respectively. The distribution of cells from each sub-cluster along pseudotime is shown at the top. (**h**) Chromatin accessibility levels of the KEs with PP/SP across the four MuSC sub-clusters. (**i**) Co-accessibility analysis between KEs and PP or SP. (**j**) Schematic illustrating the use of publicly available snATAC-seq performed in MuSCs *in vivo* from injury-induced regenerating muscles for co-accessibility analysis. (**k**) UMAP plots showing five sub-clusters of MuSCs derived from analyzing the above snATAC-seq data. (**l**) Heatmap showing the chromatin accessibility dynamics of the PP/SP and the KEs along the pseudotime trajectory inferred from the above *in vivo* snATAC-seq data. Green and purple rectangles highlight KEs with accessibility patterns co-varying with PP and SP, respectively. The distribution of cells from each sub-cluster along pseudotime is shown at the top. (**m**) Chromatin accessibility levels of the KEs and PP/SP across the five MuSC sub-clusters. (**n**) Co-accessibility analysis between the KEs and PP or SP. (**o**) 4C-seq analysis showing interactions between the indicated KEs and PP (Left) or SP (Right) in Ctrl and PP-Krab C2C12 cells. (**p**) 4C-seq analysis showing interactions between the indicated KEs and PP (Left) or SP (Right) in Ctrl and SP-Krab C2C12 cells. (**q**) Schematic showing the dual sgRNAs mediated targeted deletion of KE (KE-KO) in C2C12 myoblasts. (**r**-**u**) qRT-PCR analysis of *PP-Runx1* and *SP-Runx1* expression in the above KE1 (**r**), KE12 (**s**), KE13 (**t**), or KE22 (**u**) knockout cells. (**v**) Schematic of CRISPR/Cas9/AAV-sgRNA mediated *in vivo* knockout of KEs in MuSCs. FISCs were harvested and cultured to ASC-D1, ASC-D2 and DSC-D3 cells for detecting *PP-* and *SP-Runx1* expressions. (**w**-**z**) qRT-PCR detection of *PP-Runx1* and *SP-Runx1* expression in the above sgKE cells. Statistical significance was calculated using Student’s t-test. *p < 0.05, **p < 0.01, ***p < 0.001 and ns, no significance.

To further dissect KE-PP/SP interaction dynamics, we performed single-nucleus ATAC-seq (snATAC-seq) in FISCs, ASC-D1 and ASC-D2 (Fig. 4d). Unsupervised clustering of the quality-controlled cells (Extended Data Fig. 4a-4c) using t-distributed stochastic neighbor embedding (t-SNE) revealed distinct global chromatin accessibility profiles across the three stages (Extended Data Fig. 4d). Further clustering based on gene scores for MuSC stage-specific markers—quantified by the chromatin accessibility of each gene locus—identified four transcriptionally distinct clusters: close-to-quiescence (cQ) MuSCs, early-activating MuSCs (eA), medium-activating (mA) and late activating (lA) MuSCs (Fig. 4e). Consistent with prior results from analyzing scRNA-seq^40^, cQ cells exhibited high gene scores for quiescence-associated markers (*Pax7*, *Spry1*, *Heyl*, *Notch3*), whereas eA cells showed elevated accessibility at loci encoding activation (*Myod1*) and cell cycle related genes (*Cdk6*) (Fig. 4f). To delineate the temporal relationship between PP or SP and the KEs, we performed pseudotime trajectory analysis of chromatin accessibility (Fig. 4g and Supplementary Table 3), which partitioned the KEs into two distinct modules. One class (including KE1 and KE22), similar to PP, demonstrated significant openness early in cQ and eA cells followed by subsequent gradual closure (Fig. 4g-4h); the other class (including KE11-KE14), resembling SP, gradually opened later in mA and lA cells. Co-accessibility scoring uncovered that KE1 and KE22 openness showed a strong positive correlation with PP, while KE6, KE11-KE14 exhibited a positive correlation with SP (Fig. 4i and Extended Data Fig. 4e). To assess whether these temporal associations are conserved *in vivo*, we analyzed publicly available snATAC-seq datasets from adult mouse skeletal muscle undergoing injury-induced regeneration^41^ (Fig. 4j). Uniform manifold approximation and projection (UMAP) analysis stratified MuSCs into five sub-populations^41^: QSCs, self-renewing MuSCs (SSCs), ASCs, and two differentiating states (DSC1, DSC2) (Fig. 4k and Extended Data Fig. 4f). Pseudotime trajectory analysis of these *in vivo*-derived cells largely recapitulated the modular organization of PP/SP and associated KEs observed *in vitro* (Fig. 4l-4m and Supplementary Table 3). Specifically, KE1 and KE22 remained classified as the same class with PP, exhibiting high openness early in SSC and QSC that subsequently decreased. In contrast, KE11-KE14 displayed similar dynamic trends to SP, showing obvious accessibility later from ASC to DSC2. Moreover, co-accessibility patterns between the seven KEs and PP/SP were largely conserved (Fig. 4n and Extended Data Fig. 4g). To substantiate the functional involvement of KEs in PP/SP promoter competition, we found that silencing PP activity by dCas9-KRAB significantly reduced interactions between the majority of the seven KEs and PP, concomitantly enhanced the interactions between SP and most of the KEs (Fig. 4o). A similar reciprocal effect was observed upon repression of SP activity (Fig. 4p).

Next, we classified the seven KEs into three functional categories: 1. PP-to-SP switching KEs (KE1, KE22): initially interacting with PP to activate PP transcription during early activation, then switching their interaction preference to SP during later stages. 2. Induced SP-KEs (KE11, KE12, KE13): progressively gaining accessibility to enhance SP-driven transcription during late activation. 3. Induced dual REs (KE6 and KE14): increasing interaction with both SP and PP during activation to potentiate transcription from both promoters. To assess the functional impact of PP-to-SP switching KEs and SP-induced KEs, we deleted two from each group: KE1, KE22, KE12, KE13 in C2C12 myoblasts (Fig. 4q and Extended Data Fig. 4h). Deletion of KE1 had no significant effect on either *PP*- or *SP-Runx1* expression, likely due to enhancer redundancy (Fig. 4r). In contrast, deletion of KE12 (Fig. 4s), KE13 (Fig. 4t), or KE22 (Fig. 4u) specifically reduced *SP-Runx1* expression without affecting *PP-Runx1*, indicating their preferential role in regulating SP during the proliferative phase. To validate these findings *in vivo*, we deleted these KEs in endogenous MuSCs leveraging the CRISPR/Cas9/AAV9-sgRNA system (Fig. 4v-4z and Extended Data Fig. 4i). Consistent with their “switching” function, deletion of KE1 or KE22 reduced *PP-Runx1* expression in FISCs but had no effect on its subsequent expression (Fig. 4w and 4z, left); however, these deletions impaired *SP-Runx1* expression during later stages (Fig. 4w and 4z, right). Conversely, deletion of KE12 or KE13 specifically diminished *SP-Runx1* expression in late stage without altering *PP-Runx1* levels (Fig. 4x-4y), aligning with their induced activity to activate *SP-Runx1* expression. Collectively, through the above findings we have identified a cohort of hub KEs that play distinct roles to dynamically orchestrate *Runx1* PP/SP promoter competition during MuSC activation.

### Identification of USF1 as a key factor driving *Runx1* PP/SP promoter switch

Next, we sought to identify key protein factors mediating the E-P rewiring to orchestrate PP/SP switch. We first tested whether CTCF plays a role given emerging evidence showing CTCF binding at promoters or proximal regions determines promoter choice^42,43^. CTCF ChIP–seq indeed revealed a prominent binding peak exclusively at the PP in FISC, whereas both PP and SP exhibited CTCF occupancy in ASC-D2 (Extended Data Fig. 5a and Supplementary Table 2), which coincided with the induced expression of *SP-Runx1* during MuSC activation, suggesting a putative function for CTCF in PP/SP competition. However, when we deleted each CTCF-binding site (CBS) within the CTCF peaks flanking each promoter (two CBSs per peak) (Extended Data Fig. 5a-5c), no significant reduction of the corresponding *Runx1* isoforms was observed (Extended Data Fig. 5d), dampening our enthusiasm for pursuing CTCF. Of note, luciferase reporter assays using PP or SP as the core promoter revealed that PP activity declined when MuSC progressed from activation to proliferation, whereas SP activity increased (Fig. 5a), suggesting that promoter strength may govern PP/SP competition: the induced SP activity enables it to competitively recruit enhancers away from PP, thereby promoting *SP-Runx1* expression.

**Figure 5.**
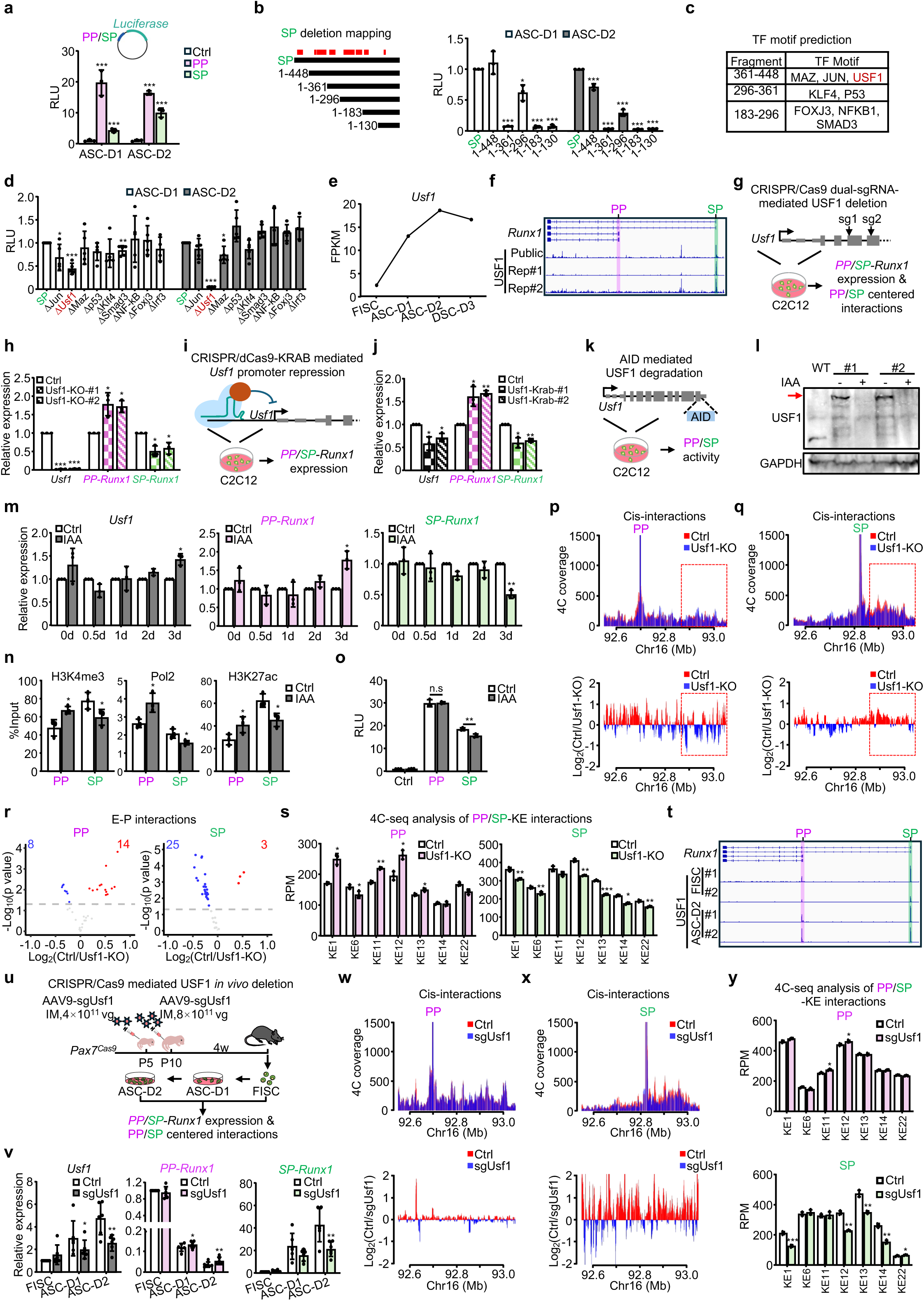
Identification of USF1 as a key factor driving *Runx1* PP/SP promoter switch. (**a**) Luciferase reporter assay measuring the promoter activity of PP and SP in ASC-D1 and ASC-D2. Diagram of the constructed reporter plasmids is shown. (**b**) Left: Schematic of SP deletion mapping. Red bars indicate regions enriched for TF binding motifs. Right: Luciferase reporter assay in ASC-D1 and ASC-D2 measuring the activity of the indicated SP fragments. (**c**) *De novo* prediction of candidate TFs with binding motifs located within the indicated SP fragments. **(d)** The above predicted TF motifs were each deleted in the reporter and the impact on SP promoter activity was detected by luciferase reporter assays in ASC-D1 and ASC-D2. (**e**) RNA-seq detection of *Usf1* expression dynamics during MuSC lineage progression. (**f**) Analyzing publicly available and in-house generated USF1 ChIP-seq in C2C12 cells showing USF1 binding profiles at the PP and SP. (**g**) Schematic showing the dual sgRNAs mediated targeted knockout of USF1 in C2C12 myoblasts. (**h**) qRT-PCR analysis of *Usf1*, *PP-Runx1*, and *SP-Runx1* expression in the USF1-KO cells. (**i**) Schematic of dCas9-KRAB-mediated repression of *Usf1* transcription in C2C12 myoblasts. (**j**) qRT-PCR analysis of *Usf1*, *PP-Runx1*, and *SP-Runx1* expression in the above Usf1-Krab cells. (**k**) Schematic of generating C2C12 cell line with inducible USF1 degradation using the AID2 system. The mAID tag was inserted at the C-terminus of the endogenous USF1 protein. (**l**) Western blot showing USF1 degradation following IAA treatment in the above USF1-AID cells. The degraded protein bands are indicated with red arrows. GAPDH was used as a loading control. (**m**) qRT-PCR detection of *Usf1* (Left), *PP-Runx1* (Middle) and *SP-Runx1* (Right) expressions in the above USF1-AID cells treated with IAA for the indicated times. (**n**) ChIP-qPCR analysis of H3K4me3 (Left), RNA Pol II (Middle), and H3K27ac (Right) occupancy at PP and SP in the above cells. (**o**) Luciferase reporter detection of PP and SP activity in the above cells (**p**-**q**) 4C-seq analysis showing the change of PP/SP centered cis-interactions in Ctrl and Usf1-KO cells. Top: Normalized 4C-seq coverage profiles showing overlaid contact signals at PP (**p**) or SP (**q**) locus in Ctrl (Red) vs. Usf1-KO (Blue). The reads are normalized to 1M total reads. Bottom: Differential cis-interactions centered on PP (**p**)- or SP (**q**) between Ctrl and Usf1-KO. The region upstream of the SP is highlighted with a red rectangle. (**r**) The number of increased (Red) or decreased (Blue) E-P interactions at PP (Left) and SP (Right) in Ctrl vs. Usf1-KO. (**s**) Normalized 4C-seq reads showing interactions between the indicated KEs and PP (Left) or SP (Right) in Ctrl and Usf1-KO C2C12 cells. (**t**) USF1 ChIP-seq profiles at PP and SP in FISC and ASC-D2. (**u**) Schematic of *in vivo* knockout of USF1 in MuSCs. AAV9 vectors encoding sgRNAs targeting *Usf1* coding regions (4 × 10¹¹ vg/mouse at P5, followed by 8 × 10¹¹ vg/mouse at P10) were injected intramuscularly into *Pax7^Cas9^* mice. Control mice received the same dose of AAV9 virus containing the pAAV9-sgRNA backbone without sgRNA insertion. FISCs were isolated four weeks later and cultured to ASC-D1 and ASC-D2 for analysis. (**v**) qRT-PCR detection of *Usf1* (Left), *PP-Runx1* (Middle) and *SP-Runx1* (Right) expressions in the above cells. (**w**-**x**) 4C-seq analysis showing the change of PP/SP centered cis-interactions in Ctrl and sgUsf1 cells at ASC-D2. Top: Normalized 4C-seq coverage profiles showing overlaid contact signals at PP (**w**) or SP (**x**) locus in Ctrl (Red) vs. sgUsf1 (Blue). The reads are normalized to 1M total reads. Bottom: Differential cis-interactions centered on PP (**w**)- or SP (**x**) between Ctrl and sgUsf1. (**y**) Normalized 4C-seq reads showing interactions between the indicated KEs and PP (Top) or SP (Bottom) in Ctrl and sgUsf1 MuSCs. Statistical significance was calculated using Student’s t-test. *p < 0.05, **p < 0.01, ***p < 0.001 and ns, no significance.

To identify candidate TFs driving AP activation, we next systematically mapped functional regions within the SP to identify potential factors driving its activation. A series of truncated luciferase reporter constructs spanning positions 1-448 bp, 1-361 bp, 1-296 bp, 1-183 bp, and 1-130 bp of SP (Fig. 5b, left) were constructed for deletion mapping; luciferase assays revealed that sequences within regions 361-448 bp, 296-361 bp, and 183-296 bp were critical for driving the reporter activity both in activation (ASC-D1) and proliferation (ASC-D2) stages (Fig. 5b, right). Motif enrichment analysis within these regions identified several candidate TFs including MAZ, JUN, USF1, KLF4, P53, FOXJ3, NFKB1 and SMAD3 (Fig. 5c). Next, each TF-binding motif was individually deleted to assess the impact on SP activity. Strikingly, only deleting the USF1-binding motif markedly reduced SP-driven reporter activity especially at the proliferation stage (ASC-D2) (Fig. 5d) when *SP-Runx1* expression was highly induced (Fig. 1c), indicating its essential role in SP activation. Consistent with this, *Usf1* expression increased during MuSC activation (Fig. 5e and Extended Data Fig. 6a), which mirrored *SP-Runx1* dynamics (Fig. 1c). Importantly, both publicly available and in-house ChIP-seq data in myoblasts showed USF binding enrichment selectively to the SP (Fig. 5f and Supplementary Table 2). To test whether USF1 regulates SP activation and promoter competition, C2C12 myoblasts with CRISPR/Cas9-mediated *Usf1* knockout were generated (Fig. 5g and Extended Data Fig. 6b-6c). *Usf1* deletion reduced *SP-Runx1* expression while concomitantly increasing *PP-Runx1* levels (Fig. 5h), supporting a role for USF1 in promoting SP activity and mediating promoter competition. Moreover, dCas9-KRAB-mediated transcriptional repression of *Usf1* (Fig. 5i) resulted in decreased *SP-Runx1* and increased *PP-Runx1* expression (Fig. 5j). Finally, inducible degradation of USF1 using the AID system resulted in decreased *SP-Runx1* and a corresponding increase in *PP-Runx1* (Fig. 5k-5m and Extended Data Fig. 6d-6f), accompanied by corresponding changes in Pol II, H3K4me3 and H3K27ac signals (Fig. 5n and Extended Data Fig. 6g). Luciferase reporter assays further substantiated these findings: USF1 depletion selectively impaired transcriptional activity of SP, without affecting PP activity (Fig. 5o and Extended Data Fig. 6h), sustaining the functions of USF1 in driving SP activation and PP/SP competition.

**Figure 6.**
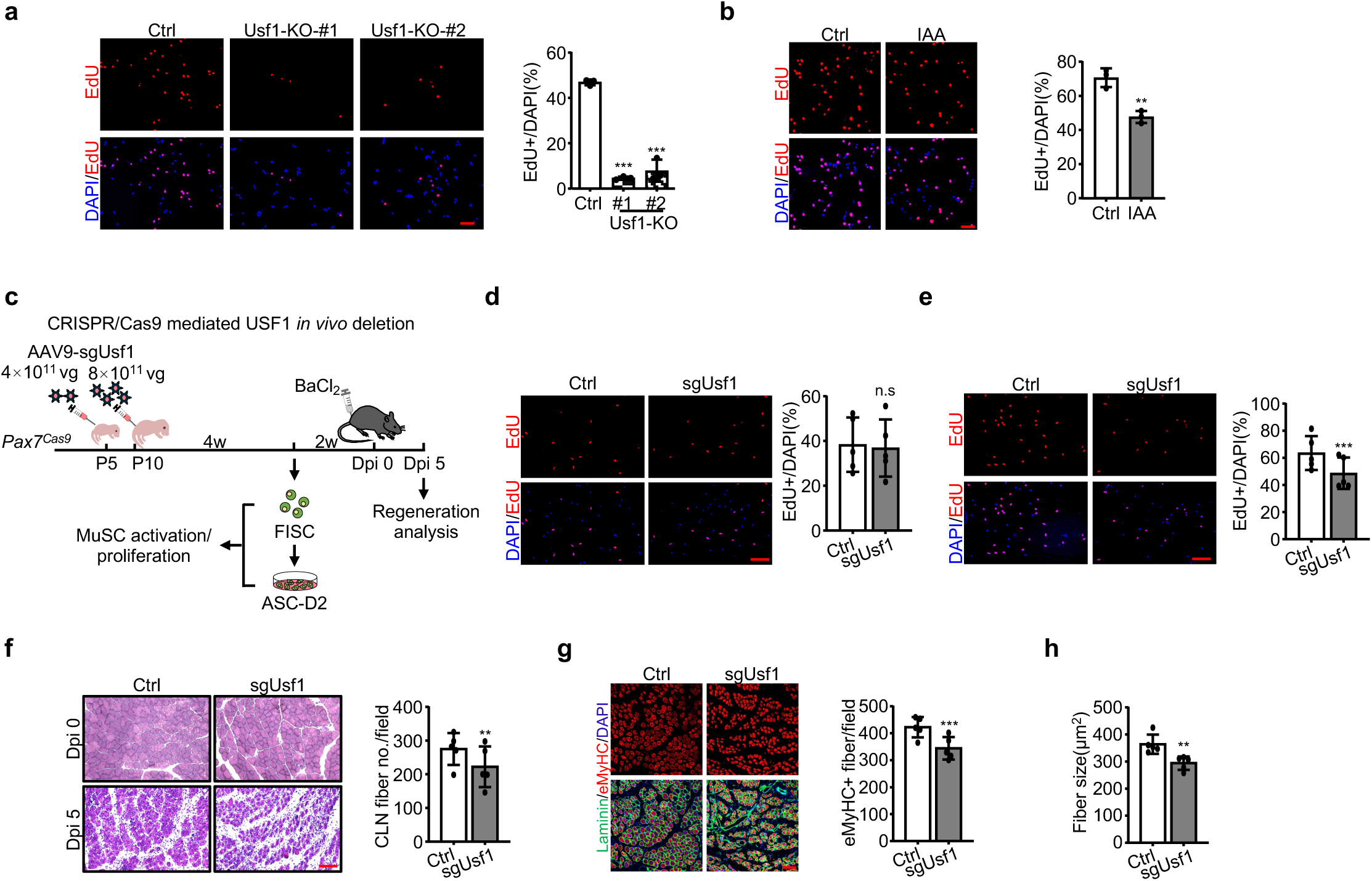
Functional involvement of USF1 in regulating MuSC activation/proliferation and muscle regeneration. (**a**) Left: EdU staining of Ctrl and Usf1-KO C2C12 cells. Right: Quantification of the percentage of EdU+ cells. Scale bar = 100 μm. (**b**) Left: EdU staining of Ctrl and IAA USF1-AID-#1 C2C12 cells. Right: Quantification of the percentage of EdU+ cells. Scale bar: 100 μm. (**c**) Schematic of *in vivo* knockout of USF1 in MuSCs to detect its effect on muscle regeneration. AAV9 vectors encoding sgRNAs targeting *Usf1* coding regions (4 × 10¹¹ vg/mouse at P5, followed by 8 × 10¹¹ vg/mouse at P10) were injected intramuscularly into *Pax7^Cas9^* mice. Control mice received the same dose of AAV9 virus containing the pAAV9-sgRNA backbone without sgRNA insertion. FISCs were harvested and cultured to ASC-D2 cells for analysis. For muscle regeneration assays, BaCl₂ was injected into the TA muscle at least six weeks post-AAV injection (Dpi 0), and samples were collected five days later (Dpi 5) for analysis. (**d**) Left: FISCs from mice injected with Ctrl or sgUsf1 virus were labeled with EdU for 24 hours to assess changes in MuSC activation. Right: Quantification of the percentage of EdU+ cells. Scale bar = 100 μm. (**e**) Left: ASCs from the above mice were labeled with EdU to evaluate changes in MuSC proliferation. Right: Quantification of the percentage of EdU+ cells. Scale bar = 100 μm. (**f**) Left: H&E staining of TA muscles at Dpi 5 from the above mice. Right: Quantification of regenerating myofibers with CLN per field. Scale bar: 100 μm. (**g**) Left: Immunostaining of eMyHC (red) and Laminin (green) on TA sections from the above mice. Right: Quantification of eMyHC+ myofibers. Scale bar: 100 μm. **(h)** Quantification of myofiber size from sections shown in (**g**). Statistical significance was calculated using Student’s t-test. **p < 0.01, ***p < 0.001 and ns, no significance.

To further dissect whether USF1 controls SP/PP competition through modulating E-P rewiring, we performed 4C-seq and found that *Usf1* deletion in the Usf1-KO cells markedly strengthened interactions between PP and upstream enhancer-rich regions (Fig. 5p), whereas diminishing cis-interactions of SP with the regions (Fig. 5q). Consistently, PP-anchored E-P interactions globally increased, concomitant with a reciprocal decrease in SP-anchored interactions (Fig. 5r). Most KEs showed enhanced interactions with PP, while their interactions with SP significantly diminished (Fig. 5s), highlighting a role for USF1 in active rewiring of the E-P interactions to participate in PP/SP competition.

Lastly, to substantiate the regulatory role of USF1 in MuSCs *in vivo*, we performed USF1 ChIP-seq in FISC and ASC-D2 (Fig. 5t and Supplementary Table 2). At both stages, robust USF1 binding was observed at SP. In contrast to the findings in C2C12 myoblasts (Fig. 5f), we also detected USF1 binding to PP in MuSCs, albeit with a lower intensity. Notably, we failed to identify canonical USF1 binding motif at PP, suggesting that USF1 may occupy this region indirectly. To further validate the regulatory role of USF1 in SP, we knocked out USF1 in endogenous MuSCs using the CRISPR/Cas9/AAV9-sgRNA system (Fig. 5u and Extended Data Fig. 6i). As expected, USF1 loss markedly suppressed the induction of *SP-Runx1* expression during MuSC activation, while the corresponding downregulation of *PP-Runx1* was partially alleviated (Fig. 5v and Extended Data Fig. 6j). Further 4C-seq showed that the changes in PP-centered cis-interactions were less pronounced compared to those observed in C2C12 myoblasts upon USF1 knockout (Fig. 5p and 5w); however, SP-centered interactions were substantially diminished (Fig. 5x). Interactions of KEs with SP were broadly reduced (Fig. 5y, bottom). Although interactions with the PP remained largely stable, we nonetheless observed a modest increase with KE11 and KE12 following USF1 knockout (Fig. 5y, top). Altogether, the above results establish USF1 as a key TF regulator that binds directly to SP to drive competitive enhancer engagement to facilitate SP activation and thus control *Runx1* isoform output.

### Functional involvement of USF1 in regulating MuSC activation/proliferation and muscle regeneration

USF1 is a ubiquitously expressed TF that plays a critical role in lipid and glucose metabolism^44^ as well as cardiometabolic health^45^ but its potential role in muscle is unknown. To demonstrate the functional relevancy of USF1 in MuSCs and regeneration, we found that both genetic deletion and inducible degradation of USF1 impaired myoblast proliferation (Fig. 6a-6b and Extended Data Fig. 7), phenocopying the effects observed upon SP deletion (Fig. 2). To examine the functional consequences of USF1 deficiency in MuSC activities and muscle regeneration *in vivo*, six weeks after intramuscular injection of AAV-sgRNA virus into the *Pax7^Cas9^*mice, BaCl₂-induced muscle injury/regeneration was assessed at Dpi 5 (Fig. 6c). USF1 knockout had minimal impact on MuSC activation (Fig. 6d) but significantly impaired MuSC proliferation (Fig. 6e), which was consistent with the phenotypes observed upon SP-RUNX1 deletion (Fig. 2f-2g). Moreover, USF1 deficiency resulted in a reduced number of newly regenerating myofibers with CLN (Fig. 6f). IF staining for eMyHC also confirmed a marked decrease in the number and size of the newly formed myofibers (Fig. 6g-6h), indicating the delay of muscle regeneration. Altogether these results demonstrate the functional relevance of USF1 in regulating MuSC activation/proliferation and muscle regeneration.

### Global identification of AP usage during MuSC activation and promoter competition as a general phenomenon

Finally, we sought to investigate whether AP usage is a general phenomenon occurring at other loci during MuSC activation. To this end, we quantified transcriptional activity at each promoter using the *proActiv* tool^3^ based on RNA-seq derived expression levels from FISC and ASC-D2 (Fig. 7a-7b). As expected, 53.09% of genes used more than one promoter (Fig. 7c). For each gene, the promoter with the highest activity was designated as “major promoter”; promoters with activity < 0.25 were classified as “inactive promoters”, while the remaining constituted “minor promoters”. In total we identified 12,183 major, 14,473 minor and 833 inactive promoters in FISC, and 12,186 major, 13,190 minor and 2,113 inactive promoters in D2 (Fig. 7d). While most promoters retained their identity between the two stages, a subset underwent switching (Fig. 7d). For example, of the 12,183 major promoters identified in FISC, 10,499 retained their status in ASC-D2, whereas 1,501 transitioned to minor promoters and a smaller subset (183) became inactive. To evaluate the accuracy of expression-based estimation of promoter activity, we found major promoters exhibited the highest H3K4me3, H3K27ac and ATAC-seq signals, whereas inactive promoters showed the lowest (Fig. 7e and Extended Data Fig. 8a-8b).

**Figure 7.**
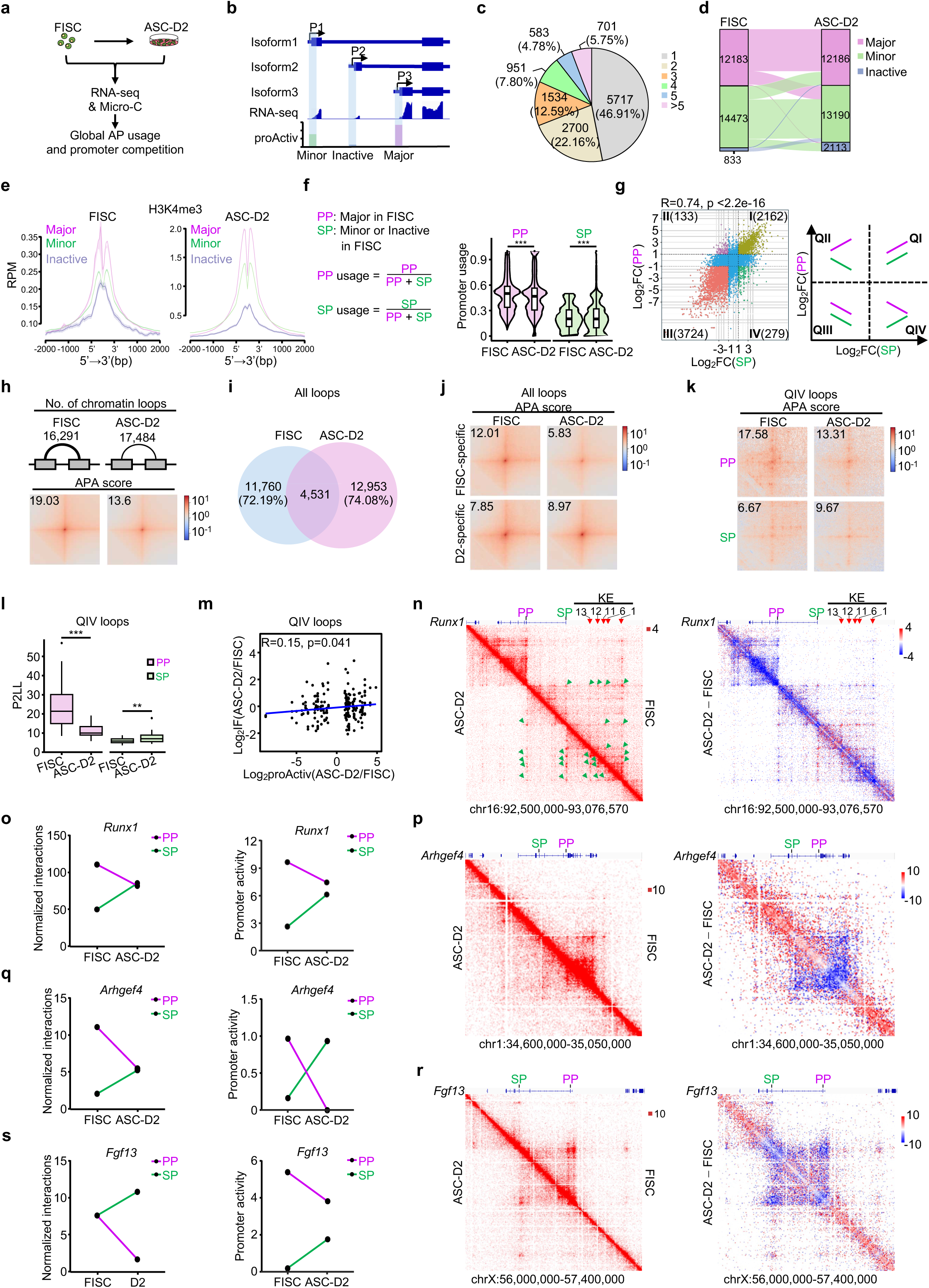
Global identification of AP usage during MuSC activation and promoter competition as a general phenomenon. (**a**) Schematic of the integrative analysis to investigate global AP usage and promoter competition during MuSC activation. (**b**) Schematic of promoter activity quantification using the *proActiv* algorithm based on RNA-seq data. For each gene, the promoter with the highest activity was designated as “major promoter”; promoters with activity < 0.25 were classified as “inactive promoters”, while the remaining constituted “minor promoters”. (**c**) Pie chart displaying the distribution of genes harboring the indicated number of promoters. (**d**) The number of promoters in each class in FISC and ASC-D2. (**e**) H3K4me3 ChIP-seq signal profiles at each promoter class in FISC (Left) and ASC-D2 (Right). Signal intensity is plotted within a ± 2 kb window centered on the transcription start site (TSS) for each promoter. (**f**) Left: Formulas used to calculate usage of primary (major, PP) and secondary (minor and inactive, SP) promoters. Right: Violin plots illustrating PP vs. SP usage in FISC and ASC-D2. (**g**) Left: Scatter plot showing classification of PP/SP pairs into four quadrants according to the promoter activity change in ASC-D2 vs. FISC. The number of PP/SP pairs in each quadrant is labeled. Correlation between changes in PP and paired SP activity is shown. Right: Schematic illustrating the trends in promoter activity changes for PP and paired SP in each quadrant. (**h**) Top: The number of identified chromatin loops in FISC and ASC-D2. Bottom: APA analysis of the above all chromatin loops. (**i**) Venn diagram showing the remodeling of chromatin loops in FISC vs. ASC-D2. (**j**) APA analysis of the FISC- or D2-specific chromatin loops. (**k**) APA analysis of chromatin loops anchored on PP and paired SP in Quadrant IV (QIV) of (**g**). (**l**) P2LL analysis of loop strength centered on the above PP /SP pairs. (**m**) Correlation between changes in promoter activity and associated loop interaction frequency (IF) for PP/SP in QIV. (**n**) Left: Micro-C contact maps at the *Runx1* locus in FISC and ASC-D2. Positions of PP and SP are marked and positions of KEs are indicated with red arrows. Dots marked by green arrows represent chromatin interactions among PP or SP and the KEs. Right: Comparison of contact frequencies at the *Runx1* locus between FISC and ASC-D2. (**o**) Left: Normalized chromatin interaction frequencies (Left) and promoter activities (right) for *Runx1* PP and SP in FISC and ASC-D2. (**p**) Left: Micro-C contact maps at the *Arhgef4* locus in FISC and ASC-D2. Positions of PP and SP are marked. Right: Comparison of contact frequencies at the *Arhgef4* locus between FISC and ASC-D2. (**q**) Left: Normalized chromatin interaction frequencies (Left) and promoter activities (right) for *Arhgef4* PP and SP in FISC and ASC-D2. (**r**) Left: Micro-C contact maps at the *Fgf13* locus in FISC and ASC-D2. Positions of PP and SP are marked and positions of KEs are indicated with red arrows. Right: Comparison of contact frequencies at the *Fgf13* locus between FISC and ASC-D2. (**s**) Left: Normalized chromatin interaction frequencies (Left) and promoter activities (right) for *Fgf13* PP and SP in FISC and ASC-D2. Statistical significance was calculated using a Student’s t-test for (**g**) and (**m**). **p < 0.01, ***p < 0.001.

**Figure 8.**
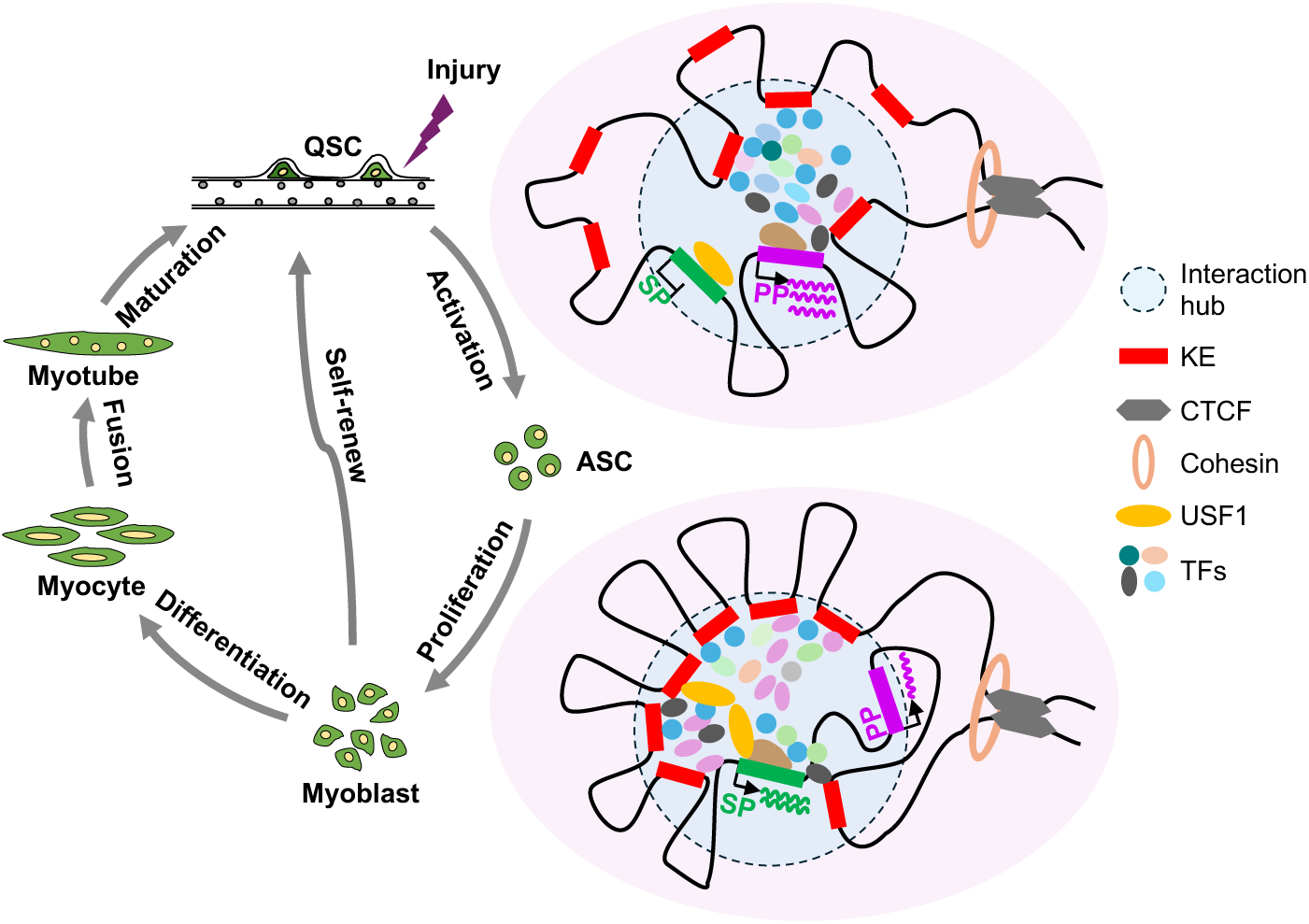
Proposed model of a promoter competition hub mediating *Runx1* PP/SP promoter switch during MuSC activation. A dynamic promoter competition hub comprising PP, SP, and a set of KEs orchestrates *Runx1* AP usage within a TAD maintained by CTCF and cohesin during MuSC activation. During the QSC-to-FISC transition, the PP is selectively activated to initiate MuSC activation. A subset of KEs, assisted by TFs, interacts with the PP within a multi-connected interaction hub, while the SP remains in a closed, transcriptionally silent state. During this phase, USF1 acts as a pioneer factor by binding to the SP to prime it for subsequent activation. During the FISC-to-ASC transition, USF1 facilitates the opening of the SP, which, together with other TFs, drives *SP-Runx1* expression to promote MuSC proliferation. Due to the competitive relationship between the two promoters, KEs that previously interacted with the PP switch their associations to the SP. Concurrently, additional KEs initially located outside the hub are recruited to interact with the SP to boost its rapid expression, while a small subset of these KEs retains interaction with the PP to maintain a basal level of *PP-Runx1* expression.

To calculate changes in promoter utilization, we defined the major promoter of each gene in FISC as the PP and minor or inactive promoters of the same gene as SP; PP or SP usage was computed as the ratio of PP or SP activity to the total promoter activity per gene^4^ (Fig. 7f, left). As a result, we observed a global decrease in PP usage and a corresponding increase in SP usage during MuSC activation (Fig.7f, right). When profiling the dynamic activity of PPs and SPs (Fig. 7g and Supplementary Table 4), we found most PP/SP pairs showed concordant changes in activity (located in Quadrants I and III), a small subset exhibited opposing trends (Quadrants II and IV). Specifically, 279 PP/SP pairs (Quadrant IV) displayed a competitive pattern analogous to the PP/SP switch in *Runx1*(Fig. 7g). Gene ontology (GO) enrichment analysis of the genes in each quadrant revealed distinct functional categories (Extended Data Fig. 8c-8f and Supplementary Table 1), suggesting the dynamic promoter usage is associated with diverse functions.

We next interrogated whether changes in promoter activity correlate with alterations in chromatin looping by performing high-resolution Micro-C in FISC and ASC-D2 (Fig. 7a). Approximately 1.4 billion valid pairs were obtained to enable visualization of chromatin loops at 1 kb resolution (Supplementary Table 5). Global looping analysis uncovered the number of identified loops increased from 16,291 in FISCs to 17,484 in ASC-D2, while aggregated peak analysis (APA) indicated a marked decrease in global loop interaction strength (19.03 vs. 13.6) (Fig. 7h). Consistent with our prior findings from analyzing Hi-C data^33^, global characterization revealed a substantial loop remodeling and 72.19% and 74.08% stage-specific loops were identified in FISCs and ASC-D2, respectively (Fig. 7i). As expected, FISC-specific loops showed a marked reduction in APA score in D2 vs. FISC (5.83 vs. 12.01), while D2-specific loops exhibited elevated APA score (8.97 vs. 7.85) (Fig. 7j). Consistently, interaction frequency decreased for FISC-specific loops and increased for D2-specific loops (Extended Data Fig. 8g).

When examining the 279 competitive PP/SP pairs in Quadrant IV (Fig. 7k), decreased PP activity indeed correlated with attenuated PP-centered interaction strength (APA: 17.58 in FISC vs. 13.31 in ASC-D2), while increased SP activity was accompanied by enhanced SP-centered looping (APA: 6.67 vs. 9.67), mirroring the pattern observed for *Runx1*. In contrast, for PP/SP pairs in other quadrants, interaction strength centered on both PPs and SPs generally diminished (Extended Data Fig. 8h-8j). These findings suggest that, at least for competitive PP/SP pairs, changes in promoter activity were positively coupled with local looping interactions. To further validate this association, we quantified loop strength centered on PP and SP in Quadrant IV using the Peak to Lower Left (P2LL) metric^46^. Again, PP activity loss correlated with diminished loop strength, whereas SP activation coincided with strengthened looping (Fig. 7l). Moreover, across all promoters in Quadrant IV, changes in promoter activity showed a modest but statistically significant positive correlation with changes in loop interaction frequency (Fig. 7m). As expected, Micro-C data analysis on *Runx1* locus recapitulated the findings from using Hi-C and 4C-seq (Fig. 3i-3l): PP-centered interactions markedly diminished from FISC to ASC-D2, while those centered on SP increased (Fig. 7n and 7o, left), paralleling their respective promoter activities (Fig. 7o, right). A close examination also detected interactions involving five KEs, including KE1, KE6, KE11, KE12, and KE13, with SP and PP (Fig. 7n). Importantly, interactions were observed between SP and PP as well as among these KEs, confirming these elements interact with each other to form a multi-connected hub. Such promoter competition model was also clearly observed on many other loci, including *Arhgef4* (Fig. 7p-7q) and known key regulators of MuSC function such as *Fgf13*^47^ (Fig. 7r-7s). Collectively, our results establish AP usage and PP/SP promoter competition as a general phenomenon during MuSC activation and chromatin loop rewiring as an underlying mechanism driving PP/SP promoter switch, thereby contributing to the precise spatiotemporal control of gene expression.

## Discussion

In this study, we leveraged *Runx1* locus as a paradigm and MuSC activation as a biological setting to dissect AP usage and the underlying regulatory mechanism. We first uncovered that the two *Runx1* promoters, PP and SP derived isoforms exhibit opposing expression dynamics during MuSC activation and fulfill distinct functional roles. Further investigations revealed that these two promoters mutually repress each other’s expression through engaging in antagonistic chromatin interactions. We then identified a set of KEs dynamically interacting with these two promoters to form a multi-connected E-P hub, orchestrating the PP/SP competition during MuSC activation. Furthermore, USF1 was identified as a critical TF that directly binds SP to modulate the promoter competition (Fig. 8). Lastly, we also found that AP usage represents a general phenomenon during MuSC activation and chromatin interaction orchestrated promoter competition governs the AP usage of a subset of genes.

### Alternative promoter usage provides an additional regulatory layer for MuSC fate transitions

Our study highlights the important regulatory role of RUNX1 protein in MuSC lineage progression during muscle regeneration. By dissecting the distinct functional contributions of these two promoters derived protein isoforms, we provide a compelling paradigm for how AP selection orchestrates MuSC lineage progression. We demonstrated a precise temporal switching between these promoters: *PP-Runx1* is rapidly induced to govern early MuSC activation but is dispensable for proliferation, whereas *SP-Runx1* expression increases subsequently to specifically drive myoblast expansion (Fig.1 and 2). This intricate regulatory arrangement highlights a critical function of AP usage in enabling a single genetic locus to uncouple and independently execute discrete, stage-specific cellular functions during dynamic cell fate transitions. Of note, this mechanism extends far beyond *Runx1*: over 50% expressed genes identified in FISC and ASC-D2 harbor more than one promoter (Fig. 7c), indicating AP usage is a general phenomenon in MuSC activation.

AP selection can provide multi-layered control over gene expression. While different promoters typically generate transcript isoforms with differing 5′-UTR lengths^1,4^—thereby fine-tuning mRNA stability and translational efficiency—it can also dramatically expand the cellular proteome repertoire. As demonstrated by our *Runx1* data, PP/SP switching produces functionally distinct protein variants tailored to specific lineage stages. This principle is mirrored in other developmental contexts; for instance, previous studies have shown that a mouse-specific retrotransposon functions as an AP to produce a truncated Cdk2ap1ΔN protein isoform, which is strictly required for cell proliferation during early embryogenesis^48^. Thus, the widespread utilization of APs represents a critical additional layer of gene regulation. By generating isoform-specific temporal expression patterns and substantially expanding proteomic functional diversity, AP usage equips MuSCs with the extraordinary regulatory flexibility required to precisely navigate the complex fate transitions.

### A dynamic promoter competition hub orchestrates *Runx1* AP usage during MuSC activation

AP usage has been known as a widespread phenomenon across diverse species, but the underlying mechanism is not well characterized. Leveraging *Runx1* locus as a paradigm, our study is the first to dissect how E-P chromatin interactions can contribute to AP usage and promoter choice. Of note, our findings highlight E-P rewiring as a novel mechanism orchestrating AP usage. We identified a multi-connected E-P hub comprising PP, SP, and multiple associated KEs. E-P hubs are emerging as key regulatory entities in coordinating gene transcription^13,18,49^. Although highly connected multiway E-P hubs containing multiple promoters and enhancers have been previously described^13,49,50^—often associated with high transcriptional output and enriched for cell identity genes—such hubs are typically thought to facilitate co-regulation, with all promoters simultaneously interacting with shared enhancers and exhibiting concordant expression^13,50^. In the case of *Runx1*, however, despite residing within a multi-connected hub, PP and SP exhibit clear competitive behavior (Fig. 3). In both stable-state proliferating C2C12 myoblasts and dynamically fate-transitioning MuSCs, inhibition or knockout of one promoter leads to clear increased activity of the other, demonstrating that this competitive relationship is not transient or context-dependent^25^.Thus, our findings indicate that promoters within multi-connected hubs are not always transcriptionally synchronized, highlighting the functional complexity of hubs. Beyond *Runx1*, through the global profiling of AP usage, we identified a subset of PPs and SPs with inversely correlated activities, accompanied by alterations in 3D interactions (Fig. 7), suggesting loop rewiring mediated promoter competition is not an isolated phenomenon. We think the competition can also apply to promoters of adjacent genes. Indeed, a competition between the *Myc* promoter and its proximal lincRNA PVT promoter was reported^51^.

Notably, the multi-connected hub encompassing *Runx1* locus is present in both FISC and ASC-D2 but SP transcription is absent in FISCs (Fig. 8), in line with the notion that some hubs exist before gene expression begins; this suggests a priming mechanism that poises the SP for rapid activation upon a signaling cue being received. Nevertheless, the hub itself is far from static; rather, it undergoes dynamic rewiring through shifting KE activity and interaction preferences (Fig. 4). We identified three classes of hub-associated KEs. Among these, the PP-to-SP switching KEs, including KE1 and KE22, display openness dynamics similar to PP, promoting its transcription in early activation stage, followed by a gradual decline in activity and a shift in interaction from PP to SP. Knockout of these KEs affects PP expression during MuSC early activation, while their impact on SP becomes evident in later stages (Fig. 4w and 4z). The induced-SP KEs, such as KE12 and KE13, however showed induced activity when MuSCs activate and interact with SP to facilitate its activation. Their knockout impacts SP expression in the later stages but has no effect on PP (Fig. 4x-4y). Such hub maintains KEs and both promoters in proximity and permits dynamic E-P interaction remodeling within the hub. As a result, the hub enables rapid redistribution of the transcriptional machinery between the PP and SP upon cell activation cue, and a swift downregulation of *PP-Runx1* and a rapid induction of *SP-Runx1*. Thus, the hub encompassing the *Runx1* locus likely represents a unique class of multi-connected E-P hubs. Instead of being transient or stage-specific^13^, these hubs function as long-lived but dynamic and adaptable regulatory platforms. Through intrinsic rewiring of E-P interactions, they enable rapid redistribution of transcriptional machinery among internal promoters to precisely coordinate stage-specific promoter expression dynamics^18^.

### USF1 binding at SP induces its activation and modulates promoter selection within the hub

Further mechanistic investigation into PP/SP competition with the hub reveals that CTCF may not be a key driver despite CTCF binding near SP and PP correlates with their expression dynamics (Extended Data Fig. 5). Of note, our findings identify USF1 as an important orchestrator that binds preferentially to the SP, shifting KEs to organize E-P connectivity within the hub, and drives isoform-specific transcription. Interestingly, USF1 binds directly to SP even in FISCs (Fig. 5t), when SP is closed and inactivated (Fig. 1b). It appears USF1 can induce epigenetic remodeling at SP (Fig. 5n), suggesting it may act as a pioneer factor priming for SP activation. Interestingly, disruption of the USF1 binding motif in SP almost completely abrogated its transcriptional activity (fig. 5d), whereas USF1 loss yielded only partial reduction (Fig. 5h, 5j and 5m), suggesting potential redundancy from other TFs such as USF2 and ATF2 with similar DNA-binding motifs. Thus, USF1 could act as a molecular switch that binds to SP and opens it for seizing KEs away from PP and induce SP transcription upon MuSC activation (Fig. 8). This is consistent with the emerging paradigm that promoter-intrinsic features—including sequence-specific motifs, GC content, and CpG island presence—can directly shape the compatibility of enhancers, thereby enabling precise and selective gene regulation^15,16^. It is worth noting that USF1 loss does not disrupt the presence of the *Runx1* hub (Fig. 5). Future endeavors will be needed to elucidate potential mechanism, for example, liquid–liquid phase separation (LLPS) that governs the *Runx1* hub assembly and maintenance.

## Methods

### Mice

All animal experiments were performed in compliance with the guidelines for the care and use of laboratory animals established by The Chinese University of Hong Kong (CUHK) and were approved by the Animal Experimentation Ethics Committee (AEEC) of CUHK. Mice were housed in the CUHK Animal Facility under standard laboratory conditions with a 12-hour light/12-hour dark cycle. The *Pax7-nGFP* mouse strain was kindly provided by Dr. Shahragim Tajbakhsh^52^. The Cre dependent *Rosa26*^Cas9-EGFP^ knockin mice (B6;129-Gt(ROSA)26Sor^tm1(CAG-cas9*-EGFP^) Fezh/J; stock number 024857) were obtained from the Jackson Laboratory. To generate Cas9 knockin mice, homozygous *Pax7^Cre^* mice were crossed with *Rosa26^Cas9-EGFP^* mice, as previously described^21,31,33,36,37^. To induce acute injury, ∼8-week-old mice received an injection of 50 μL of 1.2% BaCl₂ (w/v in H₂O) into the tibialis anterior (TA) muscle. TA muscles were harvested at designated time points for subsequent analysis.

### MuSC isolation by FACS

MuSCs were isolated by FACS following established protocols^21,39,53–55^. Briefly, hindlimb muscles from *Pax7-nGFP* or *Pax7^Cas9^* mice were dissected and digested with collagenase II (1,000 U/mL; Worthington) for 90 minutes at 37 °C. The resulting tissue suspension was washed with washing medium [Ham’s F-10 nutrient mixture (Sigma-Aldrich) supplemented with 10% heat-inactivated horse serum (HIHS; Gibco) and 1× penicillin–streptomycin (Gibco)]. A second enzymatic digestion was performed using collagenase II (1,000 U/mL) and dispase (11 U/mL) for 40 minutes at 37 °C to further dissociate MuSCs. Mononuclear cells were passed through a 40 μm cell strainer, and GFP⁺ MuSCs were sorted using a FACSAria Fusion cell sorter (BD Biosciences).

### Cell culture

Mouse C2C12 myoblasts (CRL-1772) and human HEK293FT cells (CRL-3216) were obtained from ATCC and cultured in DMEM supplemented with 10% fetal bovine serum (FBS), 100 U/mL penicillin, 100 μg/mL streptomycin and 2 mM L-glutamine at 37 °C in a humidified incubator with 5% CO₂. Isolated MuSCs were cultured in growth medium consisting of F10 medium supplemented with 20% FBS, 1% penicillin–streptomycin and 5 ng/mL basic fibroblast growth factor (bFGF) at 37 °C in 5% CO₂.

### EdU incorporation assay

EdU incorporation assay was done by adding EdU to the cell culture medium at a final concentration of 10 µM. Detection was performed using the Click-iT EdU kit (Invitrogen) following the manufacturer’s protocol.

### Plasmids

For the construction of *Runx1* SP or PP promoter luciferase reporter, a 526 bp sequence flanking the *SP-Runx1* transcription start site (TSS) or a 205 bp sequence flanking the *PP-Runx1* TSS was PCR-amplified from C2C12 genomic DNA and cloned into the XhoI and HindIII sites of the pGL3-Basic vector (Promega). To map regions critical for SP activity, sequences spanning positions 1-448 bp, 1-361 bp, 1-296 bp, 1-183 bp, and 1-130 bp of the SP promoter were PCR-amplified and cloned into the pGL3-Basic vector using XhoI and HindIII. For constructing RE luciferase reporters, individual REs were cloned downstream of the firefly luciferase gene in pGL3-SP or pGL3-PP via the BamHI and SalI sites. To generate pGL3-SP vectors lacking specific TF binding motifs, fusion PCR was performed using overlapping annealing sequences. For genomic editing in C2C12 cells, target-specific sgRNAs were selected using the CRISPOR web tool (http://crispor.tefor.net/)^56^ and cloned into the pX458-GFP plasmid (Addgene, 48138) using BbsI. For *in vivo* genomic editing, selected sgRNAs were synthesized and cloned into the AAV9-sgRNA transfer vector [AAV: ITR-U6-sgRNA(backbone)-CMV-DsRed-WPRE-hGHpA-ITR] via the SapI restriction site. Dual-sgRNA plasmids were generated by inserting a second sgRNA cassette containing the U6 promoter and guide sequence into the AAV9-sgRNA vector using XbaI and KpnI. All sgRNA sequences are provided in Supplementary Table 6.

### Luciferase reporter assay

For MuSCs, 4× 10⁴ cells were seeded per well of a 48-well plate and transfected with 250 ng of the indicated firefly luciferase reporter plasmids along with 10 ng of Renilla plasmid using Lipofectamine 3000. At 24 hours or 48 hours post-transfection, the luciferase activity was measured using the Dual-Glo Luciferase Assay system (Promega) according to the manufacturer’s guidelines. For C2C12 cells, transfections were performed in a 12-well format using 500 ng of the indicated firefly luciferase reporter plasmid and 20 ng of Renilla using Lipofectamine 3000 and luciferase activity was measured 48 hours post-transfection. The Luciferase/Renilla ratio was calculated for all samples.

### AAV9 virus production, purification, and injection

AAV9 virus was produced as described previously^36,37,57^. Briefly, HEK293FT cells were seeded in T75 flasks and transfected at 80–90% confluency with a 1:1:2 ratio of AAV9-sgRNA plasmid (5 μg), AAV9 serotype plasmid (5 μg), and pDF6 helper plasmid (10 μg) using polyethyleneimine. 24 hours after transfection, the medium was replaced with growth medium (DMEM supplemented with 10% FBS, 100 U/mL penicillin, 100 μg/mL streptomycin and 2 mM L-glutamine) and cells were cultured for an additional 48 hours. Cells were then harvested and washed twice with sterile phosphate-buffered saline (PBS). To release the virus, the cell pellet was resuspended in AAV lysis buffer (50 mM Tris-HCl, pH 8.0, and 150 mM NaCl) and subjected to three freeze–thaw cycles (liquid nitrogen/37 °C). The lysates were treated with Benzonase (Sigma-Aldrich) and MgCl₂ (final concentration, 1.6 mM) at 37 °C for 30 minutes, followed by centrifugation at 3,000 rpm for 10 minutes. The supernatant was mixed with 1/4 volume of 5× PEG/NaCl solution (40% PEG8000 (w/v), 2.5 M NaCl) and incubated overnight at 4 °C to precipitate the virus. The mixture was centrifuged at 4,000 g for 30 minutes at 4 °C, and the resulting pellet was resuspended in sterile PBS and centrifuged again at 3,000 g for 10 minutes. The supernatant was filtered through a 0.22 μm sterile filter and concentrated using a 100 kDa molecular weight cutoff filter (Millipore). The concentrate was washed three times with sterile PBS. Viral titers were determined by quantitative reverse transcription PCR (qRT–PCR) using primers targeting the CMV promoter. Primer sequences are listed in Supplementary Table 6. To knock out SP-RUNX1, PP-RUNX1, and USF1, heterozygous *Pax7^Cas9^*mice were intramuscularly injected with 4 × 10¹¹ vg of AAV9-sgRNA at P5, followed by a booster injection of 8 × 10¹¹ vg at P10. To target the SP/PP and regulatory elements KE1, KE12, KE13, and KE22, mice received 4 × 10¹¹ vg of AAV9-sgRNA pair #1 at P5 and 8 × 10¹¹ vg of pair #2 at P10. Control mice received the same doses of AAV9 virus containing the pAAV9-sgRNA backbone without sgRNA insertion. SCs were isolated four weeks after AAV administration.

### Genomic editing by CRISPR/Cas9 in C2C12 cells

The CRISPR/Cas9 system was used to delete SP/PP, KEs, CBSs, as well as to knockout SP-RUNX1, PP-RUNX1, total RUNX1, and USF1 expression in the C2C12 cell line. Briefly, pairs of pX458-GFP plasmids expressing sgRNAs flanking the targeted regions were transfected into C2C12 cells using Lipofectamine 3000 (Life Technologies). FACS was used to isolate EGFP+ cells 48 hours after transfection and these cells were then diluted in 96-well plates to obtain single-cell clones. Individual colonies were picked, validated by PCR, and subjected to Sanger DNA sequencing. Sequences of all sgRNAs and genotyping PCR primers are provided in Supplementary Table 6.

### Generation of mAID-inducible degradation in C2C12 cells

The AID2 system with improved degradation efficiency and reduced background degradation was applied to generate mAID-inducible degradation C2C12 cells^38,39^. Briefly, a stable C2C12 cell line expressing OsTIR1(F74G) was established by co-transfecting cells with the PB-OsTIR1(F74G)-neo PiggyBac expression plasmid and a transposase vector using Lipofectamine 3000. Twenty-four hours post-transfection, cells were treated with 2.5 mg/mL G418 for two weeks to select heterogeneous populations expressing OsTIR1(F74G). For inducible degradation of SP-RUNX1 and PP-RUNX1, pX458-GFP plasmids encoding sgRNAs targeting the N-terminus of each isoform near the ATG start codon were constructed. Donor plasmids containing different antibiotic resistance markers were generated via overlap PCR using ∼500 bp homology arms flanking the BSD/HygR-P2A-mAID-mCherry2 sequence (Addgene, 121180 and 121183), and cloned into the pRK5-Flag backbone via BamHI and HindIII sites. For inducible degradation of USF1, sgRNAs targeting the C-terminal region near the stop codon of each isoform were cloned into pX458-GFP. The corresponding donor constructs were assembled using ∼500 bp flanking sequences around the mAID-mCherry2-BSD/HygR cassette (Addgene, 72831 and 121194), and cloned into pRK5-Flag using BamHI and KpnI. pX458 and donor plasmids were co-transfected into the OsTIR1(F74G)-expressing C2C12 cells, followed by antibiotic selection using 10 µg/mL blasticidin (BSD) and 100 µg/mL hygromycin (Hygro) for two weeks prior to single-cell cloning. Successfully knocked-in clones were confirmed by genotyping PCR and Western blot. 1 µM 5-Ph-IAA was used to induce degradation of target protein. Sequences of all sgRNAs and genotyping PCR primers are provided in Supplementary Table 6.

### CRISPR/dCas9-KRAB mediated gene repression in C2C12 cells

CRISPR/dCas9-KRAB–mediated gene repression in C2C12 cells was performed as previously described^58^. Briefly, a stable C2C12 cell line expressing dCas9-KRAB was generated by co-transfecting cells with the dCas9-KRAB-containing PiggyBac expression plasmid (Addgene, 110822) and a transposase vector using Lipofectamine 3000. Twenty-four hours post-transfection, cells were subjected to selection with 10 µg/mL BSD for two weeks to enrich for heterogeneous populations stably expressing dCas9-KRAB. sgRNAs targeting genomic regions within 1 kb around gene TSS were designed using the CRISPR-ERA web tool (http://crispr-era.stanford.edu) and cloned into the lentiGuide-Puro vector (Addgene, 52963) via the BsmBI restriction site. For lentivirus production, HEK293FT cells were seeded in 10 cm dishes at 50–75% confluency and transfected with 10 μg lentiGuide-Puro-sgRNA, 8.5 μg psPAX2 (Addgene, 12260), and 4.07 μg pMD2.G (Addgene, 12259). Viral supernatant was collected and filtered through a 0.2 μm membrane filter (Pall Corporation, 4612) to remove cellular debris. To transduce C2C12 cells, 5 mL of viral supernatant was mixed with 5 mL of growth medium and added to cells in the presence of 8 μg/mL polybrene (Santa Cruz Biotechnology, sc-134220). After 24 hours, the viral medium was replaced with fresh growth medium containing 2.5 μg/mL puromycin (Thermo Fisher Scientific, A1113802). Selection continued until all cells in non-transduced control plates had died. Sequences of all sgRNAs are provided in Supplementary Table 6.

### ChIP-qPCR, ChIP-seq and data analysis

ChIP-qPCR and ChIP-seq were performed as previously described^33,59^. Briefly, approximately 500,000 cells were cross-linked with 1% formaldehyde for 10 minutes at room temperature and quenched with 125 mM glycine. Cells were then washed and collected by centrifugation at 700 g for 5 minutes at 4°C, flash-frozen in liquid nitrogen, and stored at −80°C. The cell pellet was resuspended in 130 μL of shearing buffer [10 mM Tris-HCl (pH 8.0), 0.1% SDS, and 1 mM EDTA supplemented with 1× protease inhibitor cocktail (PIC)] and subjected to sonication using a Covaris S220 instrument (intensity: 140 W, duty cycle: 5%, cycles per burst: 200, duration: 7 minutes). The resulting lysate was adjusted to a final concentration of 150 mM NaCl and 1% Triton X-100, then clarified by centrifugation at 20,000 g for 20 minutes at 4°C. The supernatant was incubated with pre-washed Dynabeads Protein G (Invitrogen, 10004D) for 2 hours at 4°C to reduce nonspecific binding. A 1/50 volume of the cleared lysate was reserved as Input control. The remaining lysate was incubated overnight at 4°C with 0.75 μg of the appropriate antibody. The following day, 10 μL of Dynabeads Protein G pre-washed in IP buffer [10 mM Tris-HCl (pH 8.0), 150 mM NaCl, 0.1% SDS, 1 mM EDTA, 1% Triton X-100, supplemented with 1× PIC] was added to each immunoprecipitation reaction and incubated for an additional 2 hours at 4°C. Beads were sequentially washed twice with IP buffer, twice with high-salt wash buffer [10 mM Tris-HCl (pH 8.0), 500 mM NaCl, 0.1% SDS, 1 mM EDTA, 1% Triton X-100], twice with LiCl wash buffer [10 mM Tris-HCl (pH 8.0), 250 mM LiCl, 1 mM EDTA, 0.5% NP-40, 0.5% sodium deoxycholate], and once with cold TE buffer [10 mM Tris-HCl (pH 8.0), 1 mM EDTA, 50 mM NaCl]. Immunocomplexes were eluted in ChIP elution buffer [50 mM Tris-HCl (pH 8.0), 10 mM EDTA, 1% SDS] by incubating at 65°C for 30 minutes. Cross-links were reversed by incubating the eluates at 65°C for 16 hours. For the Input sample, three volumes of ChIP elution buffer were added and processed in parallel with the IP samples. Samples were treated with RNase A (0.2 mg/mL) at 37°C for 2 hours, followed by Proteinase K treatment (0.2 mg/mL) at 55°C for 3 hours. DNA was then purified by phenol:chloroform extraction and ethanol precipitation, and finally resuspended in 1× TE buffer. For ChIP-qPCR, quantitative PCR was performed using Power SYBR Green Master Mix (Applied Biosystems) with primers specific to the target genomic loci. Primer sequences are listed in Supplementary Table 6. Enrichment was quantified as a percentage of Input recovery. For ChIP-seq, libraries were prepared using the NEBNext Ultra II DNA Library Preparation Kit (NEB, E7645S) and sequenced using an Illumina HiSeq X Ten or NovaSeq platform with 150 bp paired-end reads. Following antibodies were used: H3K4me3(), CTCF (ABclonal, A1133), USF1(Santa Cruz Biotechnology, sc-390027x), H3K27ac (Abcam, ab4729), PolII (Santa Cruz Biotechnology, sc-899).

Raw ChIP-seq reads were processed as previously described^33^. In brief, adapter sequences and low-quality bases were trimmed from the 3′ ends of the reads using Trimmomatic (v0.36)^60^, and reads shorter than 75 bp after trimming were discarded. The filtered reads were then aligned to the mouse reference genome (mm9) using Bowtie2 (v2.3.3.1)^61^. The resulting alignments were converted to BAM format using SAMtools (v1.5)^62^, and duplicate reads were removed using Picard (http://broadinstitute.github.io/picard). Peak calling was performed with MACS2 (v2.2.4)^63^ using a p-value threshold of 0.01, with the corresponding Input DNA sample used as a control background.

### Micro-C and data analysis

Micro-C for MuSCs was performed according to a published protocol^31^. Briefly, cells were first crosslinked at a density of 1 mL per million cells using 3 mM disuccinimidyl glutarate (DSG; MedChemExpress, HY-114697) for 35 minutes at room temperature, followed by the addition of 1% formaldehyde for an additional 10 minutes. Crosslinking was quenched with 0.375 M Tris (pH 7.0) for 5 minutes. Cells were pelleted by centrifugation at 1,000 g for 5 minutes at 4°C, resuspended in ice-cold PBS, and aliquoted at 1 million cells per tube. After centrifugation, cell pellets were snap-frozen in liquid nitrogen and stored at −80°C. For lysis, frozen cells were thawed on ice for 5 minutes and lysed in 0.5 mL of MB1 buffer [10 mM Tris-HCl (pH 7.5), 50 mM NaCl, 5 mM MgCl₂, 3 mM CaCl₂, 0.2% NP-40, and 1× PIC] for 20 minutes on ice. Cells were washed once with MB1 buffer and resuspended in 100 μL of the same buffer. The optimal MNase concentration for MuSCs was determined by prior titration experiments. Chromatin digestion was performed by adding 0.1 μL MNase (NEB, M0247S) followed by incubation at 37°C and 1,000 rpm for 20 minutes. Digestion was stopped by adding 8 μL of 500 mM EGTA and incubating at 65°C for 10 minutes. Cells were then washed twice with ice-cold MB2 buffer [10 mM Tris-HCl (pH 7.5), 50 mM NaCl, 10 mM MgCl₂]. The resulting MNase-digested DNA ends were polished using T4 polynucleotide kinase (NEB, M0201), followed by treatment with DNA polymerase I Klenow fragment (NEB, M0210). End-repair and labeling were performed using biotinylated dATP and dCTP (Jena Bioscience, NU-835-BIO14-S and NU-809-BIOXS) along with TTP and GTP. After washing with MB3 buffer (50 mM Tris-HCl, 10 mM MgCl₂), proximity ligation was carried out for 4 hours at room temperature using T4 DNA ligase (NEB, M0202). Dangling ends were removed by incubation with Exonuclease III (NEB, 0206) at 37°C for 15 minutes. Crosslinks were reversed by overnight incubation at 65°C. Following DNA extraction via ethanol precipitation, size selection was conducted using DNA purification beads to enrich ligated fragments of approximately 230 bp. Biotin-labeled fragments were then isolated using 10 μL of Dynabeads MyOne Streptavidin C1 beads (Invitrogen 65,001), and sequencing libraries were prepared using the NEBNext Ultra II DNA Library Preparation Kit. Libraries were sequenced on the Illumina NovaSeq platform with 150 bp paired-end reads.

The raw Micro-C data was mapped to the mouse reference genome (mm9) using microcket^64^ with default parameters at resolution of 10 kb, 5 kb, 2.5 kb, and 1 kb. The output .hic file from microcket was used for downstream analyses. Cool files were generated with cooler^65^ to facilitate the comparison of interaction differences. Chromatin loops were called using mustache^66^ with parameters -pt 0.01 -p 12 at resolution of 1 kb, 2.5 kb, 5 kb, and 10 kb. Annotation of loop anchor with enhancer, promoter, and CTCF features was performed by intersecting the loop anchor with ChIP-seq peaks of H3K27ac, H3K4me3, and CTCF, respectively. Aggregate peak analysis (APA) was conducted by using the coolpuppy^67^.

### 4C-seq and data analysis

4C-seq was performed as described previously^68,69^. In brief, approximately two million cells were cross-linked with 2% (v/v) formaldehyde for 10 minutes at room temperature, and quenched with 0.125 M glycine. Cells were lysed in ice-cold lysis buffer [50 mM Tris–HCl (pH 7.5), 150 mM NaCl, 5 mM EDTA, 0.5% NP-40, 1% Triton X-100, 1× PIC] for 20 minutes on ice, then centrifuged at 700 g for 5 minutes. Pelleted nuclei were washed once with 400 μl 1.2× Dpn II buffer (10× buffer diluted in H₂O), resuspended in 500 μl 1.2× Dpn II buffer, and supplemented with 15 μl 10% SDS. After incubation at 37°C for 1 hour with shaking (900 rpm), 75 μl 20% Triton X-100 was added to quench SDS, followed by 1 hour incubation at 37°C (900 rpm). For complete digestion, 200 U DpnII (NEB, R0543) was added and incubated at 37°C (900 rpm) for 4 hours. A further 200 U DpnII was added for overnight digestion at 37°C; 100 U DpnII was supplemented the next day and incubated for 24 hours to enhance efficiency. DpnII was inactivated at 65°C for 20 minutes, and samples cooled to room temperature. Ligation was performed by adding 5.7 mL H₂O, 700 μl 10× NEB T4 DNA Ligase buffer, and 3,350 U NEB T4 ligase (NEB, M0202), followed by overnight swirling at room temperature. Samples were treated with 300 μg proteinase K (Invitrogen, AM2548) at 65°C overnight, then 300 μg RNase A (Thermo Scientific, EN0531) at 37°C for 45 minutes. DNA was purified by phenol/chloroform/isoamyl alcohol extraction and ethanol precipitation, then resuspended in 150 μl 10 mM Tris–HCl (pH 7.5). Purified DNA was mixed with 50 μl rCutSmart buffer, 295 μl H₂O, and 50 U NlaIII (NEB, R0125), and incubated overnight at 37°C with shaking (500 rpm). NlaIII was inactivated at 65°C for 25 min. A second ligation was performed with 12.1 mL H₂O, 1.4 mL 10× NEB T4 DNA Ligase buffer, and 6,700 U T4 ligase overnight at room temperature. DNA was purified (phenol/chloroform/isoamyl alcohol extraction, ethanol precipitation), resuspended in 250 μl Tris-HCl (pH 7.5), and further purified with the QIAquick PCR Purification Kit (QIAGEN, 28104). For library preparation, 800 ng DNA was used as template for 20 PCR cycles with Phanta Master Mix (Vazyme, P51101). PCR products were purified with the NucleoSpin Gel and PCR Clean-up Kit (MACHEREY-NAGEL, 740609). 4C libraries were prepared using the NEBNext Ultra II DNA Library Preparation Kit and sequenced on the Illumina NovaSeq platform with 150 bp paired-end reads. Primer sequences for 4C-seq are listed in Supplementary Table 6.

4C-seq reads were processed using pipe4C^69^. Raw reads containing the 4C primer sequence were retained, adapter-trimmed, and aligned to the mouse reference genome (mm9) using Bowtie2 (v2.4.2)^61^. Only reads uniquely mapped to DpnII restriction fragment ends were retained for analysis. Final coverage tracks were generated as bigWig files, normalized to reads per million (RPM) based on the total number of mappable reads per sample, and used for downstream visualization. Overlay of 4C-seq data of different conditions is processed by the in-house script. To identify the differential interactions between different conditions, we extracted the normalized interaction frequency from 4C-seq data, and then used the DESeq2 to identify the differential interactions. For the differential enhancer interactions, we overlapped the differential interactions with H3K27ac ChIP-seq peaks.

### snATAC-seq and data analysis

snATAC-seq was performed based on a previously described protocol^41,70^ with modifications. In brief, 400,0000 cells were pelleted and resuspended in 500 ul cold ATAC-Resuspension Buffer (RSB, 10 mM Tris-HCl, pH 7.4, 10 mM NaCl, 3 mM MgCl_2_) containing 0.1% NP-40, 0.1% Tween-20, and 0.01% Digitonin. After 5 minutes of incubation on ice, the nuclei were pelleted (1000 × g, 5 min, 4 °C), and washed with 500 μL cold ATAC-RSB containing 0.1% Tween-20 but NO NP-40 or digitonin. Nuclei were then resuspended in 1.2 ml 1.2×TD Buffer (2×TD buffer, 20 mM Tris-HCl pH 7.6, 10 mM 1M MgCl_2_, 20% Dimethyl Formamide) containing 0.2% Tween-20, and 0.01% Digitonin, then distributed to a 96 well plate containing 2 ul barcoded Tn5 in each well, 10 ul per well. Tagmentation was performed at 55 °C for 30 minutes with 200 rpm shaking. Reactions were quenched by adding 10 ul 40 mM EDTA to each well, followed by a 15 minutes incubation at 37 °C. Tagmented nuclei were pooled together and pelleted. Nuclei were then resuspended in 1.2 ml 10 mM Tris-HCl, pH 7.4, 10mM NaCl buffer and diluted to 20 nuclei/10 μL, then distributed into 96 well plate containing 2.5 ul barcoded i5/i7 primers in each well. Fragment release and library amplification were performed by adding 0.04% SDS (65 °C, 10 minutes), followed by neutralization with 0.4% Triton X-100, and PCR amplification using NEB Q5 High-Fidelity 2× Master Mix (72 °C, 10 minutes;18 cycles: 98 °C 30 s, 60 °C 30 s, 72 °C 30s). Unique dual-index combinations were used across four 96-well plates per replicate. Libraries were pooled, purified using MinElute columns followed by SPRI bead cleanups (1× Ampure XP), and eluted in EB buffer. Final libraries were quantified by Qubit (dsDNA HS Assay) and sequenced on the Illumina NovaSeq platform with 150 bp paired-end reads.

Raw snATAC-seq sequencing reads were demultiplexed using Cutadapt (v4.0+)^71^ and aligned to the mm9 mouse reference genome with SnapTools (v1.5+)^72^. Aligned BAM files were converted to the SNAP format and subsequently transformed into sorted fragment files using the snap-pre and dump-fragment utilities, respectively. Quality control and downstream analysis were performed using ArchR (v1.0.2)^73^. Arrow files were generated per sample, retaining only nuclei with ≥ 1,000 unique fragments and transcription start site (TSS) enrichment ≥ 8. Doublets were identified and removed using ArchR’s default doublet-scoring and filtering parameters. Dimensionality reduction was carried out using iterative latent semantic indexing (iterative LSI) via addIterativeLSI. Batch effects across samples were corrected using ArchR’s built-in Harmony integration (addHarmony) with default settings. Nuclei from all samples were jointly clustered using addClusters (resolution = 1, default), and two-dimensional visualization was performed by UMAP (addUMAP, default parameters). For nuclei derived from mononuclear cells, dimensionality reduction, sample and batch correction, and clustering were repeated using the same functions as above (default parameters). Two-dimensional projection of the reduced dimensions was further performed with ArchR’s addUMAP function, using a minimum distance of 0.3. Following UMAP projection, cell types were annotated by assigning each cluster to the cell type corresponding to the maximum chromatin accessibility of cell-type-specific marker genes. Trajectory construction was carried out using ArchR’s AddTrajectory function, guided by pre-identified clusters, biological time points, and chromatin accessibility patterns of genes associated with quiescence and muscle differentiation. Co-accessibility was performed by using the ArchR’s getCoAccessibility function.

### Targeted deep sequencing

Target deep sequencing was conducted as previously described^36,74^. Briefly, genomic DNAs from AAV9-sgRNA infected MuSCs were amplified using Q5 High-Fidelity 2× Master Mix (NEB). The PCR products were purified with VAHTS DNA Clean Beads (Vazyme, N411-01) and subsequently subjected to library preparation using the NEBNext Ultra II DNA Library Preparation Kit. Libraries containing barcodes were pooled and sequenced on the Illumina NovaSeq platform with 150 bp paired-end reads. For data analysis, the CRISPResso2^75^ was employed to calculate indel occurrences. The primers used for Deep-seq are listed in Supplementary Table 6.

### RNA isolation, qRT-PCR, RNA-seq and data analysis

Total RNA was extracted by TRIzol reagent (Invitrogen) according to the manufacturer’s instructions. cDNA was synthesized using the HiScript III 1st Strand cDNA Synthesis Kit (Vazyme, R312-01). Quantitative real-time PCR was performed on a LightCycler 480 Instrument II (Roche Life Science) with Luna Universal qPCR Master Mix (NEB, M3003L). For MuSCs, Ywhaz or Hprt1 were used for normalization. For C2C12 cells, Gapdh and 18s were used for normalization. Primer sequences are provided in Supplementary Table 6. For poly(A)+ mRNA-seq, total RNA underwent poly(A) selection (Ambion, AM61006) followed by library preparation using the NEBNext Ultra II RNA Library Prep Kit (NEB, E7770S). Libraries were sequenced as 150 bp paired-end reads on an Illumina NovaSeq platform.

Raw RNA-seq reads were processed as described in our previously publications^31,33^. Adapters and low-quality bases were trimmed from the 3′ ends, and reads <75 bp were discarded. High-quality reads were aligned to the mouse reference genome (mm9) using STAR^76^, and transcript abundances were quantified in fragments per kilobase per million mapped reads (FPKM) with Cufflinks (v2.2.1). Promoter activity was calculated using *proActiv* based on mRNA-seq according to previous publications^3^. The identification of major, minor, and inactive promoters was performed using the default parameters of *proActiv*. Promoters classified as major in FISC cells were designated as the PPs, while all others were defined as SPs. To categorize PP/SP pairs based on their dynamic activity changes during the transition from FISC to ASC-D2, we calculated the fold change in promoter activity, applying a cutoff value of 2. The pairs were classified into four distinct types: Co-upregulated PP/SP Pairs (Quadrant I in Fig. 7g): both primary and secondary promoter activities increased simultaneously; Divergent PP/SP Pairs (Quadrant II in Fig. 7g): PP activity increased while SP activity decreased; Co-downregulated PP/SP Pairs (Quadrant III in Fig. 7g): both major and alternative promoter activities decreased simultaneously; Competitive PP/SP Pairs (Quadrant IV in Fig. 7g): PP activity decreased while SP activity increased.

### Immunoblotting, immunofluorescence, and immunohistochemistry

Total cell extracts for Western blot analysis were prepared as previously described^33,59^. The following primary antibodies were used: MYOD1 (Dako, M3512, 1:2000), PAX7 (Developmental Studies Hybridoma Bank, 1:1000), MYOGENIN (Santa Cruz Biotechnology, sc-576, 1:1000), Histone H3 (Santa Cruz Biotechnology, sc-517576, 1:3000), RUNX1 (Santa Cruz Biotechnology, sc-365644, 1:1000), USF1 (Santa Cruz Biotechnology, sc-390027, 1:1000), and GAPDH (Santa Cruz Biotechnology, sc-137179, 1:2000). Hematoxylin and eosin (H&E) staining of frozen muscle cryosections was performed using standard protocol. For immunofluorescence, sections were stained with the following antibodies: Laminin (Sigma, L9393, 1:800) and anti-embryonic myosin heavy chain (eMyHC; Sigma, 1:200). Fluorescent images were acquired using a fluorescence microscope (Leica).

## Data availability

All sequencing data generated in this study have been deposited in Gene Expression Omnibus (GEO) database under the accession code GSE328933. All used datasets from other publications are summarized in Supplementary Table 7.

## Acknowledgments

Author contributions

Conceived and designed the experiments: H.W., H.S. and L.H. Methodology: H.W., H.S., L.H. and Z.W. Performed the experiments: L.H., Y.Q., Z.W. Q.Z. and L.H. Analyzed the data: Q.S. Wrote the paper: H.W., L.H., Q.S. and Y.Q. Reviewed and edited the manuscript: H.W., H.S. and L.H.

## Supplementary Materials

### List of Extended Data Figures

**Extended Data Figure 1.**
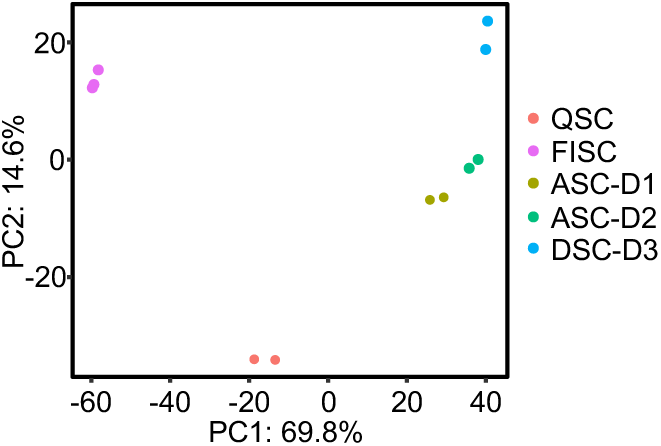
Principal component analysis (PCA) plot of RNA-seq data for MuSCs during *in vitro* lineage progression.

**Extended Data Figure 2.**
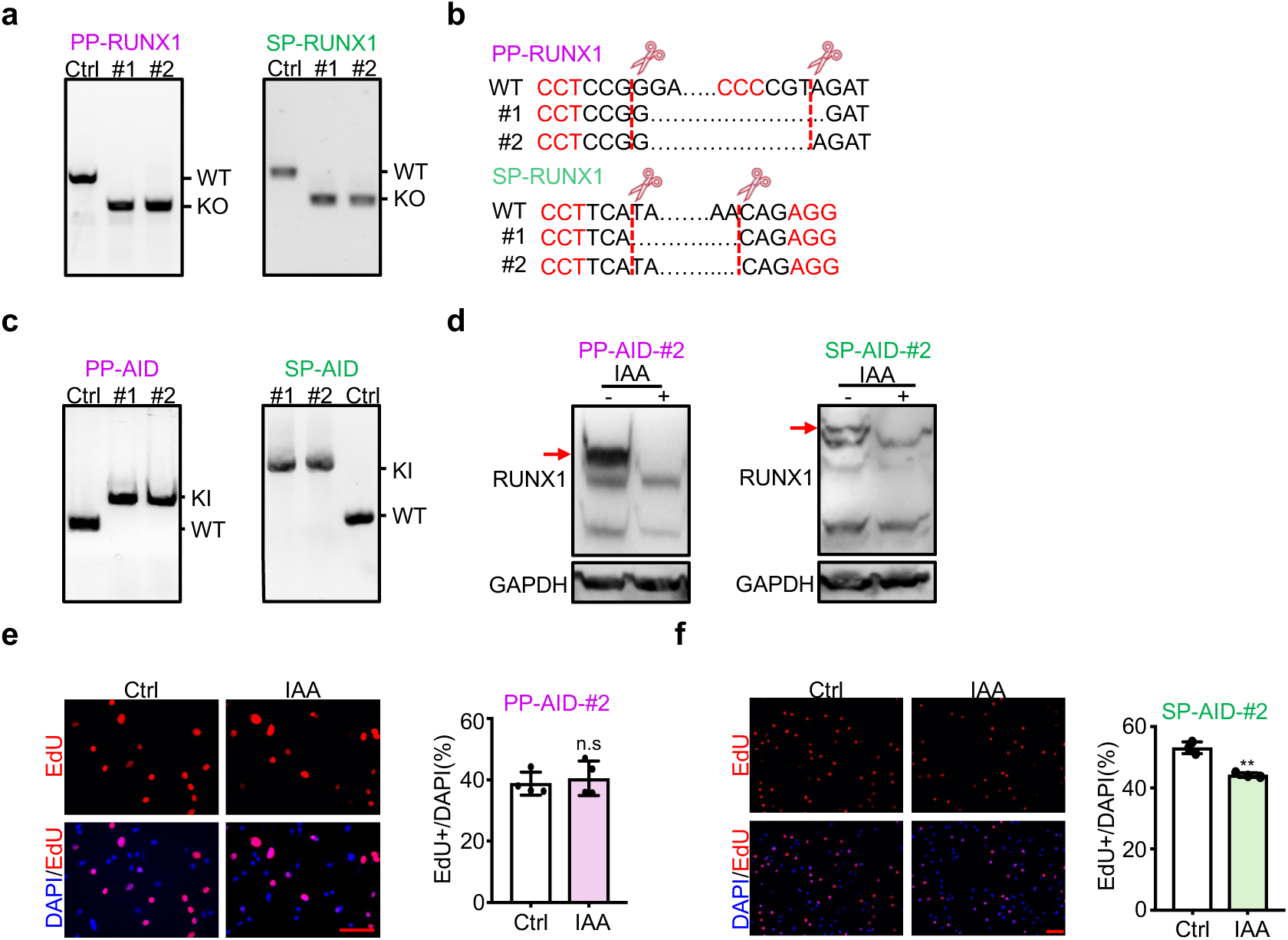
Distinct roles of PP- and SP-derived isoforms in MuSC activation/proliferation. (**a**) PCR validation of two independent C2C12 clones with *PP*- or *SP*-*Runx1* knockout (KO). PCR amplification was performed using genomic DNA from unedited control (Ctrl) and KO clones. WT and KO indicate the expected sizes of the wild-type and deleted alleles, respectively. (**b**) Sanger sequencing confirming the deletions shown in (**a**). (**c**) PCR validation of mAID knock-in (KI) to generate PP-RUNX1-AID and SP-RUNX1-AID C2C12 cell lines, with two independent clones for each line. PCR amplification was performed using genomic DNA from unedited Ctrl and KI clones. WT and KI indicate the expected sizes of the wild-type and knock-in alleles, respectively. (**d**) Western blot showing degradation of PP-RUNX1 and SP-RUNX1 following treatment of PP-AID-#2 (Left) and SP-AID-#2 (Right) clones with 5-Ph-IAA (IAA). The degraded protein bands are indicated with red arrows. GAPDH was used as a loading control. (**e**) Left: EdU staining was conducted in the above PP-AID-#2 cells after IAA treatment induced PP-RUNX1 degradation. Right: Quantification of EdU+ cells. Scale bar: 100 μm. (**f**) Left: EdU staining was conducted in the above SP-AID-#2 cells after IAA treatment induced SP-RUNX1 degradation. Right: Quantification of EdU+ cells. Scale bar: 100 μm. Statistical significance was calculated using Student’s t-test. **p < 0.01 and ns, no significance.

**Extended Data Figure 3.**
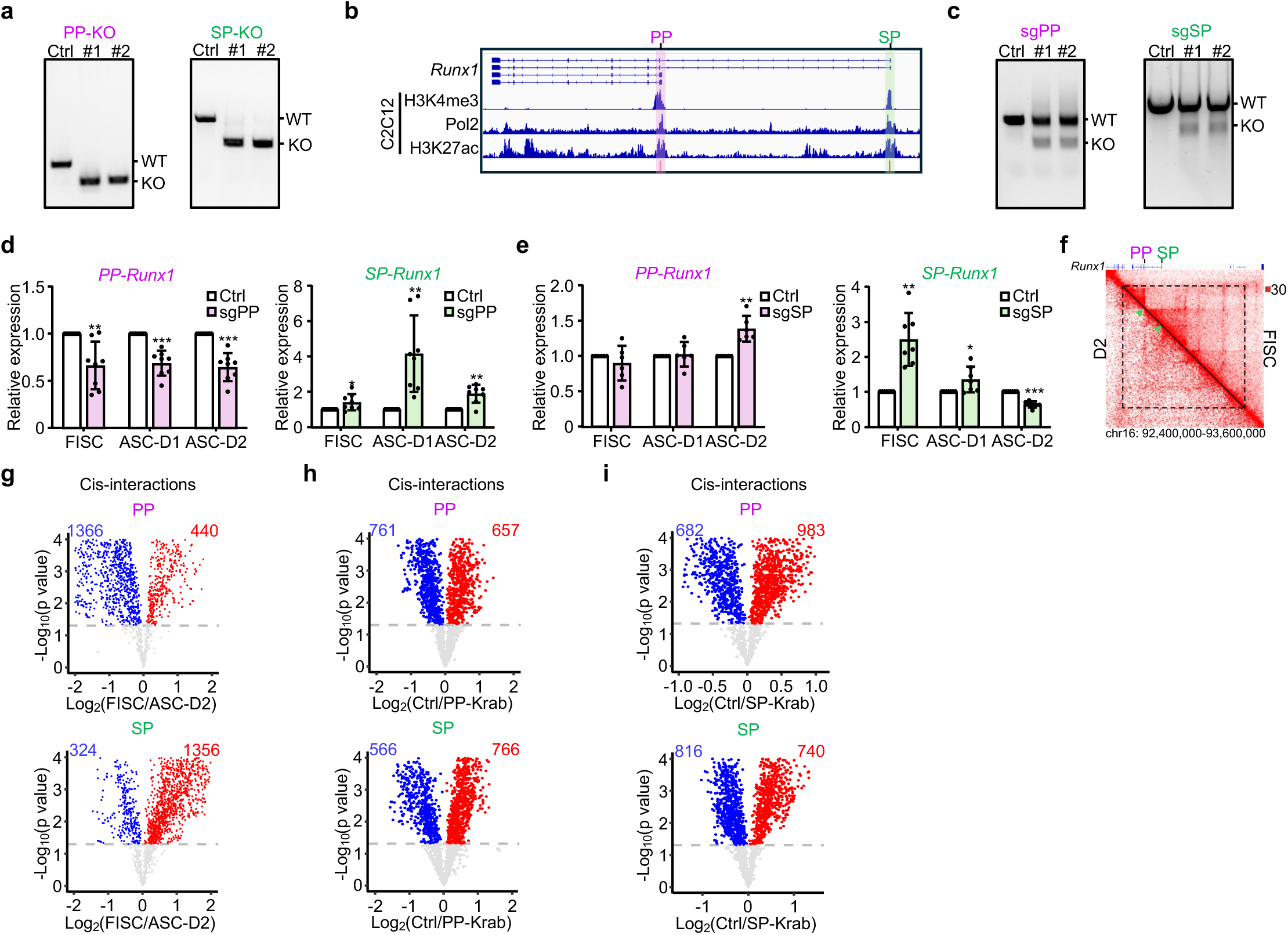
*Runx1* PP/SP promoter competition orchestrated by E-P interaction rewiring in a multi-connected E-P hub. (**a**) PCR validation of two independent C2C12 clones with PP or SP knockout. PCR amplification was performed using genomic DNA from unedited Ctrl and KO clones. WT and KO indicate the expected sizes of the wild-type and deleted alleles, respectively. (**b**) ChIP-seq profiles for H3K4me3, Pol II and H3K27ac at PP and SP in C2C12 cells. Regions for ChIP-qPCR detection are marked as red bars. (**c**) PCR validation of PP or SP knockout in MuSCs. PCR amplification was performed using genomic DNA from MuSCs isolated from two pairs of Ctrl or sgPP/SP mice. WT and KO indicate the expected sizes of the wild-type and deleted alleles, respectively. (**d**) qRT-PCR analysis showing *PP-* (Left) and *SP-Runx1* (Right) expression levels as fold change in FISC, ASC-D1 and ASC-D2 from sgPP mice. (**e**) qRT-PCR analysis showing *PP-* (Left) and *SP-Runx1* (Right) expression levels as fold change in FISC, ASC-D1 and ASC-D2 from sgSP mice. (**f**) Heatmap of our previously published Hi-C data showing contact maps at genomic regions encompassing the *Runx1* locus in FISCs and ASC-D2. The TAD harboring *Runx1* is indicated by a dashed triangular. The locations of PP and SP in the heatmap are marked with green arrows. (**g**) The number of increased (Red) or decreased (Blue) cis-interactions at PP (Top) and SP (Bottom) in FISC vs. ASC-D2. (**h**) The number of increased (Red) or decreased (Blue) cis-interactions at PP (Top) and SP (Bottom) in Ctrl vs. PP-Krab. (**i**) The number of increased (Red) or decreased (Blue) cis-interactions at PP (Top) and SP (Bottom) in Ctrl vs. SP-Krab. Statistical significance was calculated using Student’s t-test. *p < 0.05, **p < 0.01, ***p < 0.001.

**Extended Data Figure 4.**
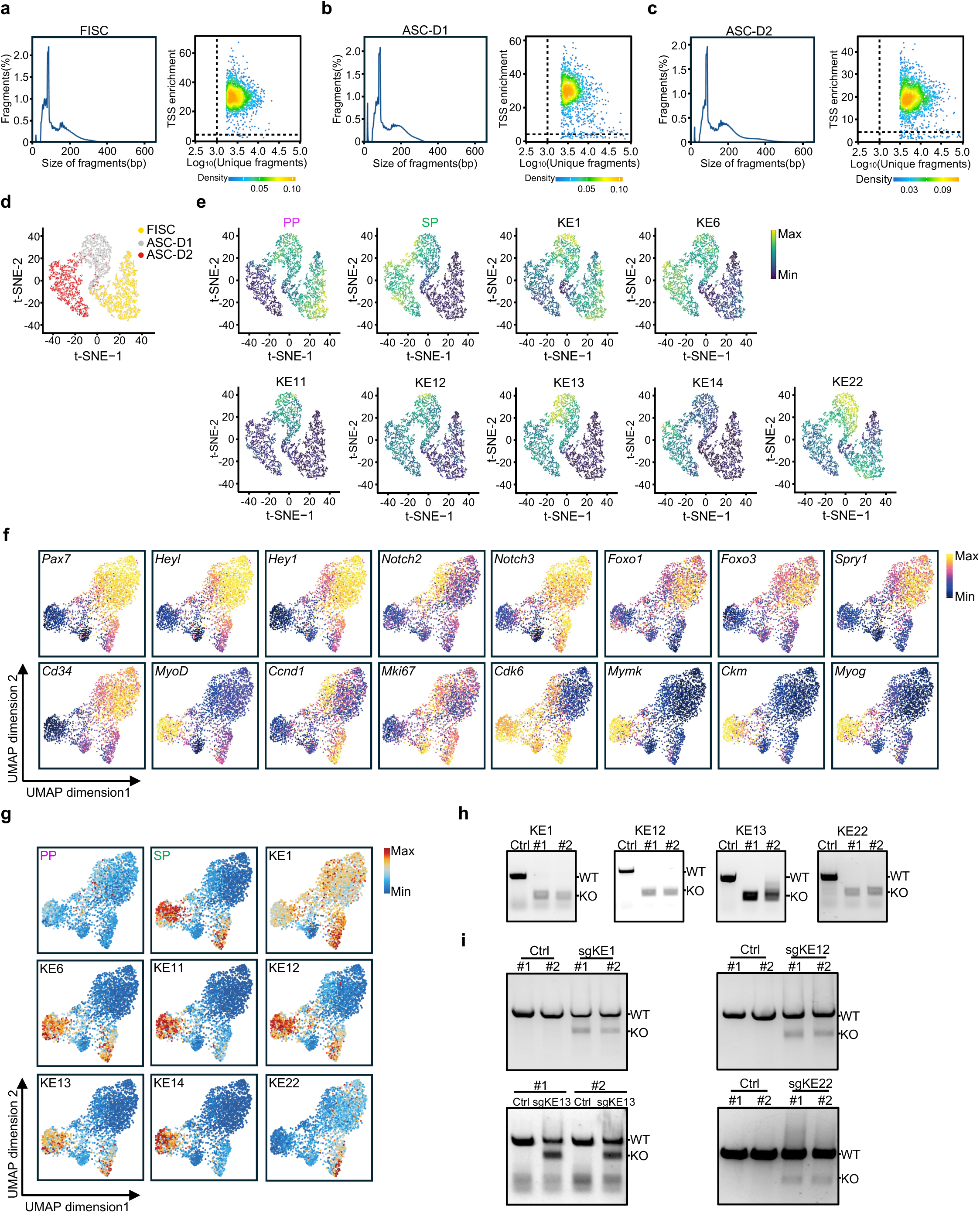
Identification of regulatory hub elements orchestrating *Runx1* PP/SP competition. (**a**-**c**) Left: Distribution of snATAC-seq fragment size in FISC (**a**), ASC-D1 (**b**) and ASC-D2 (**c**). Right: Scatter plot displaying TSS enrichment score and unique fragment number of each cell. Dashed lines indicate the filtering threshold for nuclei (Unique fragment number ≥ 10,000, TSS enrichment score ≥ 8). (**d**) t-SNE projection showing the distribution of cells in FISC, ASC-D1 and ASC-D2. (**e**) t-SNE plots displaying the accessibility of PP, SP, and KEs across the four sub-clusters (cQ, eA, mA and lA) identified based on snATAC-seq *in vitro*. (**f**) UMAP plots exhibiting the indicated gene scores across the five sub-clusters (QSCs, SSC, ASC, DSC1 and DSC2) identified based on publicly available snATAC-seq performed in MuSCs from injury-induced regenerating muscles. (**g**) UMAP plots displaying the accessibility of PP, SP, and KEs across the five sub-clusters. (**h**) PCR validation of two independent C2C12 clones with KE1, KE12, KE13 and KE22 knockout. PCR amplification was performed using genomic DNA from unedited Ctrl and KO clones. WT and KO indicate the expected sizes of the wild-type and deleted alleles, respectively. (**i**) MuSCs were isolated from Ctrl or sgKE1/KE12/KE13/KE22 mice and genomic PCR analysis was performed to test the cleavage efficiency.

**Extended Data Figure 5.**
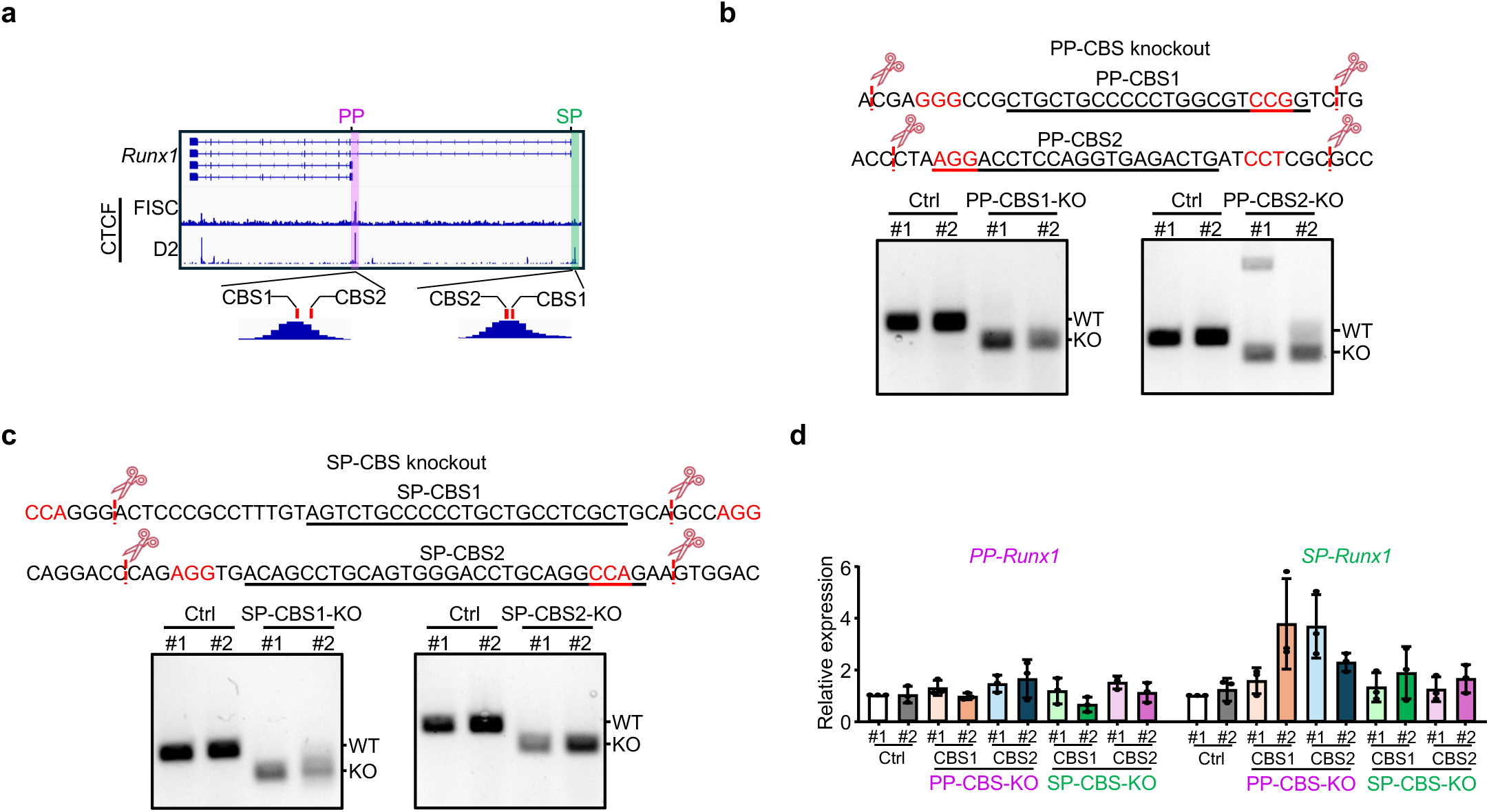
CTCF does not play a role in *Runx1* PP/SP competition. (**a**) CTCF ChIP-seq profiles at the PP and SP in FISC and ASC-D2. Two CBSs were identified at the binding peaks for each promoter. (**b**) Upper: Schematic showing the design of dual sgRNAs for targeted knockout of each CBS at the CTCF binding peaks of PP in C2C12 cells. The sequence of each CBS is displayed by underlining. Red dashed lines indicate CRISPR/Cas9 cleavage sites. Lower: PCR validation of two independent C2C12 clones with CBS knockout. PCR amplification was performed using genomic DNA from unedited Ctrl and PP-CBS-KO clones. WT and KO indicate the expected sizes of the wild-type and deleted alleles, respectively. (**c**) Upper: Schematic showing the design of dual sgRNAs for targeted knockout of each CBS at the CTCF binding peaks of SP in C2C12 cells. The sequence of each CBS is displayed by underlining. Red dashed lines indicate CRISPR/Cas9 cleavage sites. Lower: PCR validation of two independent C2C12 clones with CBS knockout. PCR amplification was performed using genomic DNA from unedited Ctrl and SP-CBS-KO clones. WT and KO indicate the expected sizes of the wild-type and deleted alleles, respectively. (**d**) qRT-PCR analysis of *PP-Runx1* and *SP-Runx1* expression in the above generated CBS KO cells.

**Extended Data Figure 6.**
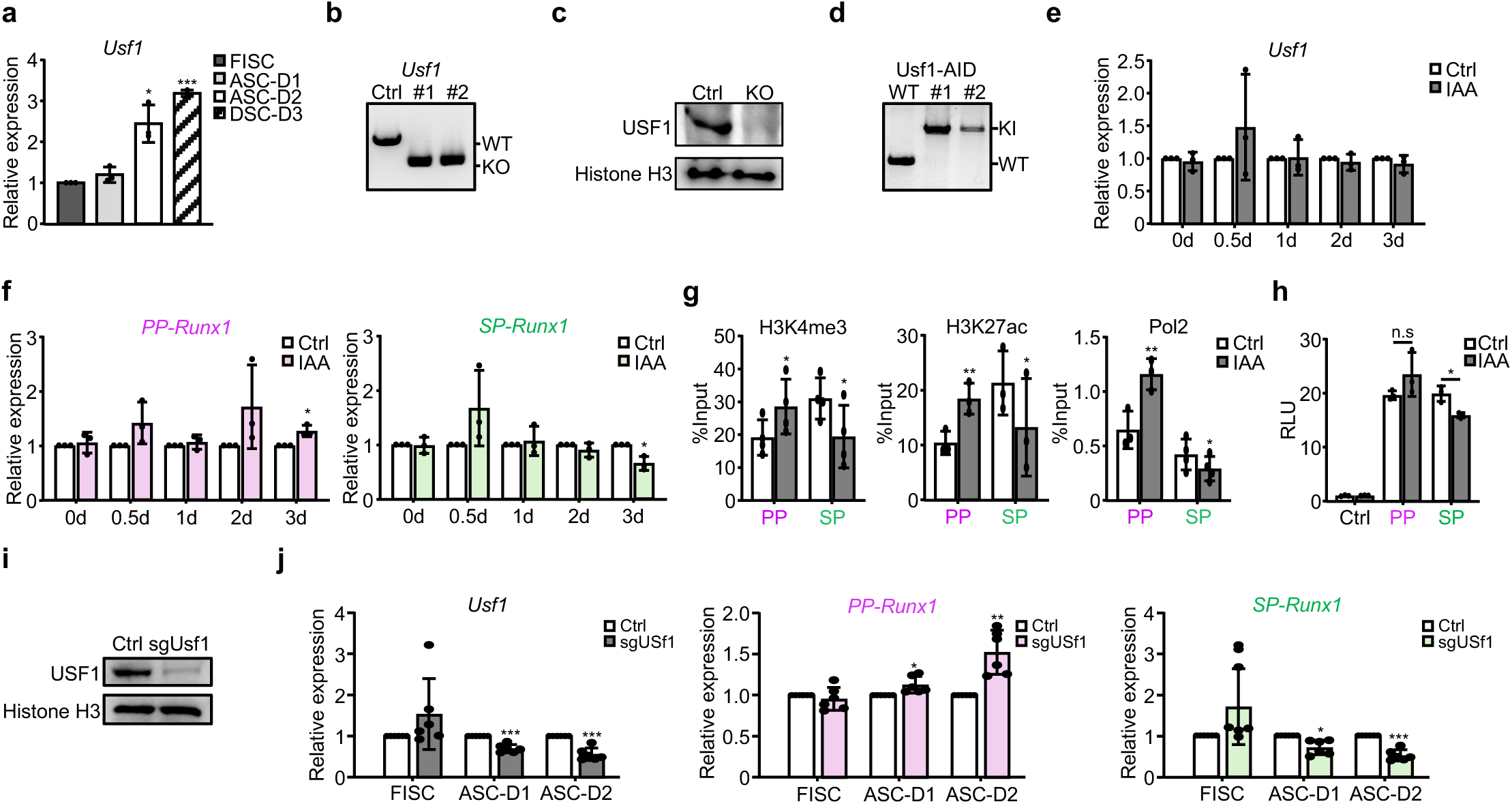
Identification of USF1 as a key factor driving *Runx1* PP/SP promoter switch. (**a**) qRT-PCR analysis showing the expression dynamics of *Usf1* during MuSC lineage progression. (**b**) PCR validation of two independent C2C12 clones with *Usf1* knockout. PCR amplification was performed using genomic DNA from unedited Ctrl and KO clones. WT and KO indicate the expected sizes of the wild-type and deleted alleles, respectively. (**c**) Western blot of USF1 protein expression in Ctrl and Usf1-KO cells. Histone H3 was used as a loading control. (**d**) PCR validation of mAID KI to generate two independent USF1-AID C2C12 cell clones. PCR amplification was performed using genomic DNA from unedited Ctrl and KI clones. WT and KI indicate the expected sizes of the wild-type and knock-in alleles, respectively. (**e**-**f**) qRT-PCR analysis to detect *Usf1* (**e**), *PP-Runx1* (**f**, Left) and *SP-Runx1* (**f**, Right) expression in the above generated USF1-AID-#2 clone treated with IAA for the indicated times. (**g**) ChIP-qPCR analysis of H3K4me3 (Left), RNA Pol II (Middle), and H3K27ac (Right) occupancy at the PP and SP in the above USF1-AID-#2 cells. (**h**) Luciferase reporter assay showing changes in PP and SP promoter activity upon induced degradation of USF1 in the above USF1-AID-#2 cells. (**i**) Western blot analysis of USF1 expression in MuSCs isolated from USF1 knockout mice (sgUsf1) compared to controls. Histone H3 served as a loading control. (**j**) qRT-PCR analysis showing *Usf1*(Left), *PP-* (Middle) and *SP-Runx1* (Right) expression levels as fold change in FISC, ASC-D1 and ASC-D2 from sgUSF1 vs. Ctrl mice. Statistical significance was calculated using Student’s t-test. *p < 0.05, **p < 0.01, ***p < 0.001 and ns, no significance.

**Extended Data Figure 7.**
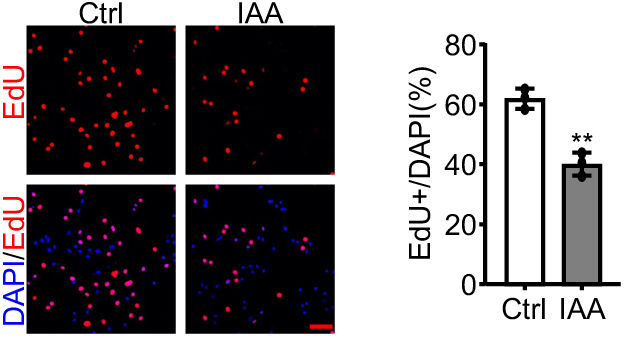
Functional involvement of USF1 in regulating MuSC activation/proliferation and muscle regeneration. Left: EdU staining was conducted in IAA-treated USF1-AID-#2 cells. Right: Quantification of EdU+ cells. Scale bar: 100 μm. Statistical significance was calculated using Student’s t-test. **p < 0.01.

**Extended Data Figure 8.**
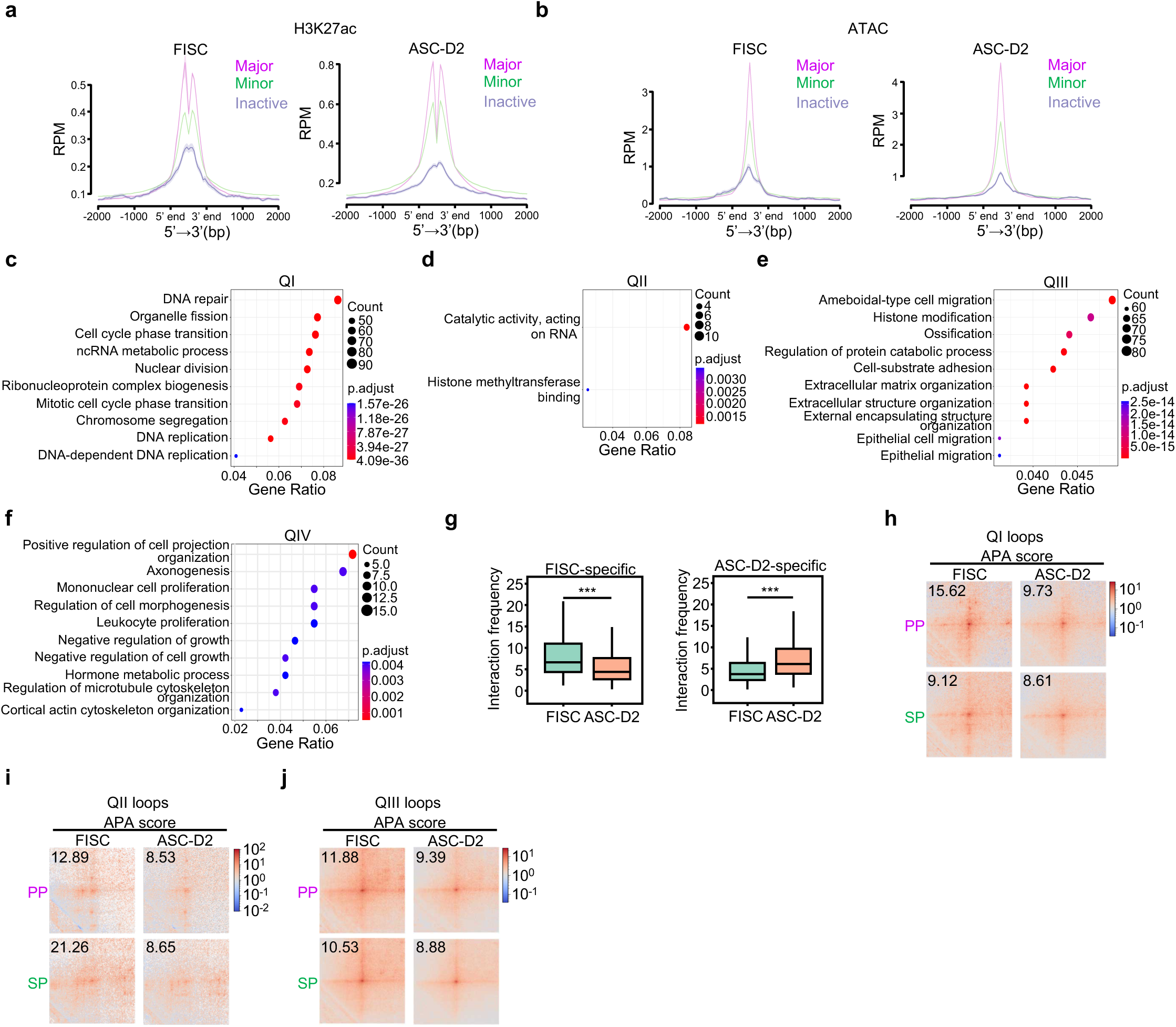
Global identification of AP usage during MuSC activation and promoter competition as a general phenomenon. (**a**-**b**) H3K27ac ChIP-seq (**a**) and ATAC-seq (**b**) signal profiles at each promoter class in FISC (Left) and ASC-D2 (Right). Signal intensity is plotted within a ± 2 kb window centered on the TSS for each promoter. (**c**-**f**) Bubble plots showing enriched GO terms associated with genes in Quadrant I (**c**), II (**d**), III (**e**), and IV (**f**) of Fig. 7g. Bubble size indicates the number of genes enriched per term and bubble color indicates the statistical significance of enrichment. (**g**) Boxplots showing interaction strengths of FISC- (Left) and D2-specific (Right) chromatin loops. (**h**-**j**) APA analysis of chromatin loops centered on PP and paired SP in Quadrant I (**h**), II (**i**) and III (**j**) of Fig. 7g. Statistical significance was calculated using Student’s t-test. ***p < 0.001.

### List of Supplementary Tables

Supplementary Table 1. RNA-seq data analysis.

Supplementary Table 2. ChIP-seq data analysis.

Supplementary Table 3. Single-nucleus ATAC-seq data analysis.

Supplementary Table 4. Differential promoter activity of PP/SP in FISC and ASC-D2.

Supplementary Table 5. Micro-C data analysis.

Supplementary Table 6. Sequences of oligos used in the study.

Supplementary Table 7. Summary of all public datasets used in the study.

